# Bridging the gap between movement data and connectivity analysis using the time-explicit Step Selection Function (tSSF)

**DOI:** 10.1101/2023.05.29.542702

**Authors:** Denis Valle, Nina Attias, Joshua A. Cullen, Mevin B. Hooten, Aline Giroux, Luiz Gustavo R. Oliveira-Santos, Arnaud L. J. Desbiez, Robert J. Fletcher

**Author notes:** Co-first authors (Denis Valle, Nina Attias). To be submitted to Movement Ecology as a “Methodology” article.

## Abstract

**Background:** Understanding how to connect habitat remnants to facilitate the movement of species is a critical task in an increasingly fragmented world impacted by human activities. The identification of dispersal routes and corridors through connectivity analysis requires measures of landscape resistance but there has been no consensus on how to calculate resistance from habitat characteristics, potentially leading to very different connectivity outcomes.

**Methods:** We propose a new model called the time-explicit step selection function (tSSF) that can be directly used for connectivity analysis in the context of the spatial absorbing Markov chain (SAMC) framework without requiring arbitrary transformations. The tSSF model combines a time model with a standard selection function and can provide complementary information regarding how animals use landscapes by separately assessing the drivers of time to traverse the landscape and the drivers of habitat selection. These models are illustrated using GPS-tracking data from giant anteaters (*Myrmecophaga tridactyla*) in the Pantanal wetlands of Brazil.

**Results:** The time model revealed that the fastest movements tended to occur between 8 pm and 5 am, suggesting a crepuscular/nocturnal behavior. Giant anteaters moved faster over wetlands while moving much slower over forests and savannas, in comparison to grasslands. We found that wetlands were consistently avoided whereas forest and savannas tended to be selected. Importantly, this model revealed that selection for forest increased with temperature, suggesting that forests may act as important thermal shelters when temperatures are high. Finally, the tSSF results can be used to simulate movement and connectivity within a fragmented landscape, revealing that giant anteaters will often not use the shortest-distance path to the destination patch (because that would require traversing a wetland, an avoided habitat) and that approximately 90% of the individuals will have reached the destination patch after 49 days.

**Conclusions:** The approach proposed here can be used to gain a better understanding of how landscape features are perceived by individuals through the decomposition of movement patterns into a time and a habitat selection component. This approach can also help bridge the gap between movement-based models and connectivity analysis, enabling the generation of time-explicit results.

## 1. Background

Land-use change is the major driver of biodiversity loss in terrestrial and freshwater ecosystems across the world (Díaz et al. 2019), causing the loss of approximately 15% of the vertebrate species richness in terrestrial landscapes (Semenchuk et al. 2022). The resulting habitat loss and fragmentation has been a central theme for conservation biologists (Harrison and Bruna 1999, Haddad et al. 2015), particularly regarding how to connect habitat remnants to facilitate the movement of species in the presence of climate change (Heller and Zavaleta 2009). Identifying wildlife dispersal routes and potential corridors through connectivity analysis typically requires the quantification of landscape resistance (Zeller et al. 2012, Hanks and Hooten 2013, Fletcher Jr. et al. 2016). Landscape resistance is often measured as a function of proxies of habitat quality, such as the estimated presence probability derived from species distribution models (e.g., Lehnen et al. 2021) or derived from habitat selection models (e.g., Zeller et al. 2016). However, an important problem with this type of connectivity analysis is that the equations commonly used to transform these proxies of habitat quality into resistance are arbitrary. For example, resistance has been often assumed to be the inverse of the predicted probability of presence (Zeller et al. 2016), but other transformations have also been applied (e.g., LaPoint et al. 2013, Lehnen et al. 2021, Iezzi et al. 2022). Furthermore, in contrast to focusing on resistance, some authors have argued that wildlife corridors should be based on areas in which animals move faster and in a directed fashion (i.e., exhibit transit behavior) (LaPoint et al. 2013, Abrahms et al. 2016) instead of areas with higher habitat quality.

In this article, we suggest that both time (or its reciprocal, speed) and habitat selection strength are distinct axes that together can help improve understanding of dispersal and connectivity across the landscape (Box 1) (see also Kuefler et al. 2010, Dickie et al. 2020). Importantly, we develop a novel habitat selection model that explicitly represents these two processes, enabling a better understanding of resource selection, and that generates a probabilistic metric of habitat selection that can be used in connectivity analysis without requiring arbitrary transformations.

Several methods have previously been developed to estimate habitat selection, with different approaches being used to characterize habitat availability for individuals or populations during a specific time frame (Hooten 2017, Northrup et al. 2022). For example, researchers often sample random locations within the boundaries of an individual’s home range to characterize habitat availability in certain types of resource selection functions (RSFs) (Manly et al. 2002). However, habitat availability for a moving individual changes through time and depends on its current location, movement capacity, and perceptual range. To incorporate this changing process in habitat availability, step selection functions (SSFs) have been increasingly used because they enable the characterization of the available habitat for each step that individuals make as they move across the landscape (Fortin et al. 2005, Forester et al. 2009). However, despite its utility, SSF does not distinguish between selection strength and how landscape characteristics influence motion capacity, potentially resulting in biased resource selection estimates. More recently, an approach called Integrated Step Selection Analysis (iSSA; Avgar et al. 2016) has been developed to account for how habitat characteristics are related both to movement and selection processes. Unfortunately, iSSA has parameter identifiability issues and can generate non-sensical parameter values for the movement kernel (see details in the Discussion section). The approach we propose avoids the problems associated with ISSA while enabling us to disentangle the effects of resource selection from landscape permeability to movement. Critically, we show how this approach can generate probabilities that can be directly used for connectivity analysis without requiring arbitrary transformation. We provide an example by modeling empirical movement data of giant anteaters (*Myrmecophaga tridactyla*) in the floodable savannas of Brazil.

### Box 1. Conceptual framework

We posit that, to understand a species’ ecology and plan its conservation, it is critical to take into account both selection strength and time to traverse the landscape. Relying just on selection strength, while ignoring time, limits the understanding of animal resource use. For example, a selected habitat might be selected for displacement (often resulting in faster movements and shorter time in that area; upper left quadrant in Fig. 1) or for resource use, such as foraging and shelter (often resulting in slower movements and longer time in the area; upper right quadrant in Fig. 1; Zollner and Lima 1999, Avgar et al. 2013). Similarly, a habitat type might be avoided because it presents a high mortality risk (often resulting in faster movements and shorter time; lower left quadrant in Fig. 1) or because it is a physical barrier to movement (often resulting in slower movements and longer time; lower right quadrant in Fig. 1; Prokopenko et al. 2017, Dickie et al. 2020, Bastille-Rousseau and Wittemyer 2021).

**Figure 1.**
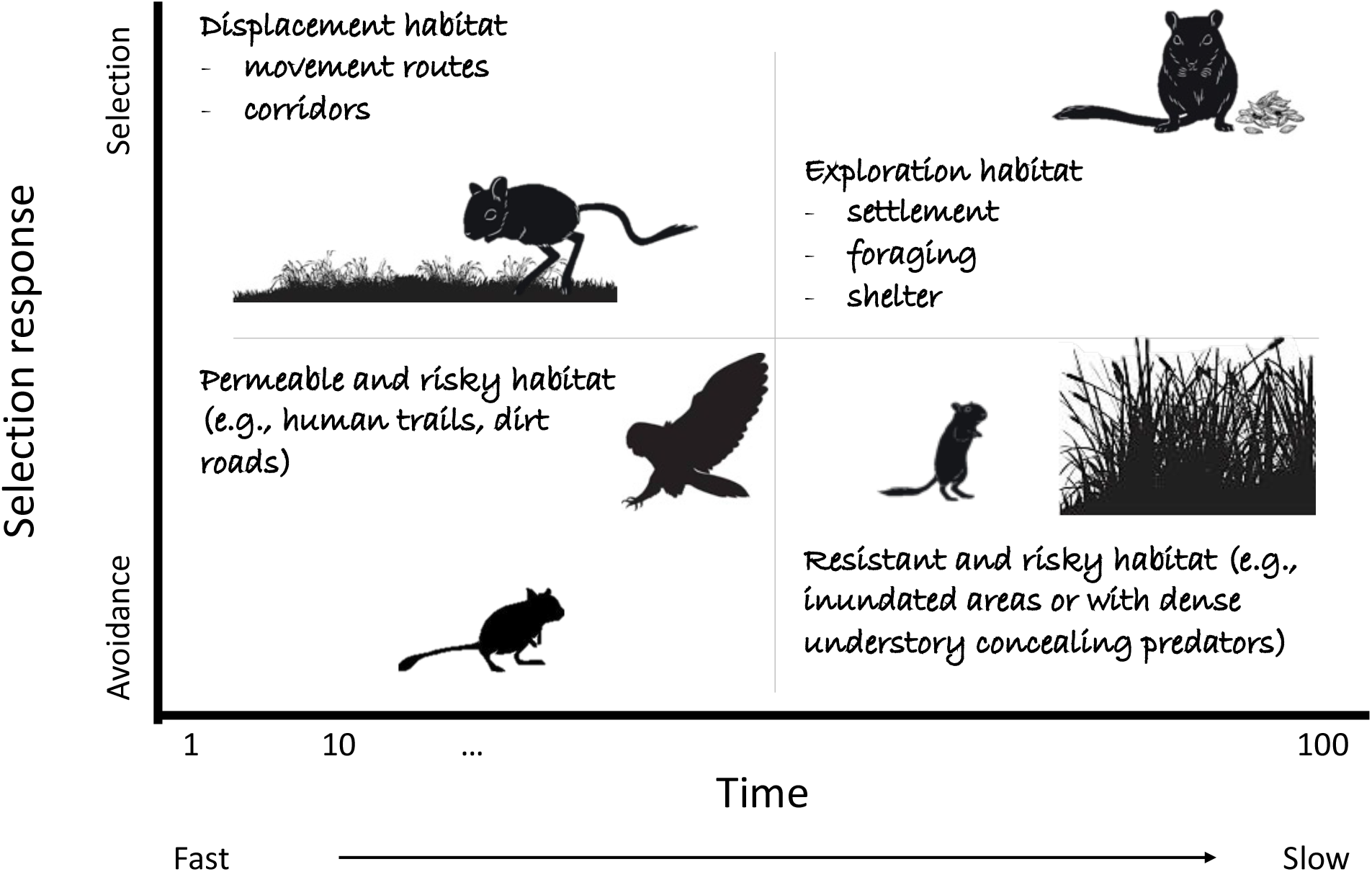
Time to traverse the landscape and selection strength as two important axes for characterizing the landscape from the perspective of a species.

Only accounting for the time taken to traverse the landscape while ignoring selection strength can also be problematic (Dickie et al. 2020). For example, residence time (i.e., the time an individual spends in a given area once it is reached) has been used as an indicator of high-quality resource areas used by animals (e.g., herbivore foraging patches or carnivore kill site; Searle et al. 2005, Van Moorter et al. 2016). However, increased time within a particular type of landscape could simply indicate greater biomechanical resistance to movement (e.g., due to greater number of obstacles or steeper slopes) (Avgar et al. 2013). Time taken to traverse a landscape can also be related to perceptual range (e.g., visually oriented small mammals tend to increase their perceptual range in habitats with low vegetation height, which allows for faster and more directed movement; Prevedello et al. 2010) and memory (e.g., prior knowledge of resource location can result in faster and more directed movement towards a given type of habitat; Fagan et al. 2013, Oliveira-Santos et al. 2016).

Examining time and selection strength as separate axes of movement is an important first step towards distinguishing between different motivations for movement. For example, we illustrate in Fig. 1 how fast movement and selection can characterize a displacement habitat (upper left quadrant) whereas slow movement and selection might be the hallmark of resource exploration habitat (upper right quadrant). Similarly, fast movement associated with avoidance might indicate permeable but risky habitat (lower left quadrant) whereas slow movement and avoidance suggest resistant and risky habitat (lower right quadrant).

In short, both selection strength and time required to traverse the landscape provide complementary insights to determine whether a landscape characteristic is perceived as a movement corridor, a source of foraging and shelter, or a source of risk, with important implications for connectivity.

## 2. Material and methods

### 2.1. Linking habitat selection models with connectivity analysis

Most of the prominent methods used for connectivity rely on estimates of cost of movement in the form of resistance surfaces (Fletcher Jr. et al. 2016, Kumar and Cushman 2022). In this article, however, we focus on parameterizing a permeability/conductance matrix (instead of a resistance surface) because this enables us to decompose movement patterns into a time and a habitat selection component.

Each cell in the permeability matrix contains the probability of choosing at a particular pixel *j* given time constraint Δ*t* and initial pixel *i* (i.e., *p*(*P*_*t*+Δ*t*_ = *j*|Δ*t*, *P*_*t*_ = *i*)). Using Bayes theorem, this probability is given by:

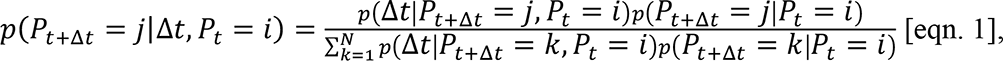

where *N* is the number of pixels in the landscape (*i*. *e*., *P*_*t*+Δ*t*_ ∈ {1, …, *N*}). Note that the time component *p*(Δ*t*|*P*_*t*+Δ*t*_ = *j*, *P*_*t*_ = *i*) quantifies the likelihood that Δ*t* will be required to reach *P*_*t*+Δ*t*_ = *j* from *P*_*t*_ = *i*, whereas the selection component *p*(*P*_*t*+Δ*t*_ = *j*|*P*_*t*_ = *i*) quantifies selection strength for pixel *j* given initial pixel *i* regardless of time constraints. This expression is similar to the equations commonly used for habitat selection models. For example, if *p*(Δ*t*|*P*_*t*+Δ*t*_ = *j*, *P*_*t*_ = *i*) is a constant for all *N* pixels, then this quantity cancels out in the numerator and denominator and this expression becomes identical to those used in standard SSF and RSF.

Although a permeability matrix is not commonly used for connectivity analysis, it is a key part of a powerful framework that has recently been proposed called the spatial absorbing Markov chain (SAMC) (Fletcher et al. 2019) and, for this reason, we rely on SAMC to link habitat selection models to connectivity analysis. This framework is based on random walk theory and captures the initiation and termination of movement, how the environment alters movement behavior, and how these processes can impact demographic rates. Aside from the permeability matrix, denoted by **Q**, SAMC may also require information on the initial distribution of a species (**Ψ**) and information on factors that may terminate movement (e.g., mortality risk from roads, settlement) (**R**). Depending on the application, all or only subsets of these components might need to be considered. Importantly, unlike other common connectivity models such as least-cost analysis or circuit-theoretic models (McRae et al. 2008, Etherington 2016), SAMC can provide time-explicit solutions in addition to long-term analytical solutions for multiple connectivity metrics.

The time-explicit nature of the SAMC allows us to directly relate our movement model results to connectivity analysis. More specifically, we develop a model to explicitly estimate *p*(Δ*t*|*P*_*t*+Δ*t*_ = *j*, *P*_*t*_ = *i*) (onwards simply “time model”) by assuming that Δ*t* is the time interval between GPS fixes. The results from this model are then used together with a selection function for *p*(*P*_*t*+Δ*t*_ = *j*|*P*_*t*_ = *i*), yielding the time-explicit Step Selection Function (tSSF). As described in Box 1, this decomposition of movement into a time component and a selection component can improve the understanding of how animals use the landscape and disperse to new areas. Below, we start by first describing the time model to then describe the tSSF.

### 2.2. Time model

We illustrate the main concepts underlying the time model using a simple hypothetical example (Fig. 2). Fig. 2a depicts the path taken by a hypothetical individual in a given step (i.e., the path defined by two consecutive GPS fixes) where step length is 6 pixels and the animal takes 7 minutes overall to traverse these pixels. The first three pixels are comprised of grasslands and the animal takes 0.8, 0.6 and 0.7 minutes to traverse these pixels whereas the next three pixels are traversed much more slowly (i.e., 1.6, 1.7, and 1.6 minutes per pixel) because they are forested pixels. Importantly, only the time taken to traverse all 6 pixels is known when using location data (i.e., the time taken to traverse individual pixels is latent and therefore has to be estimated).

**Fig. 2.**
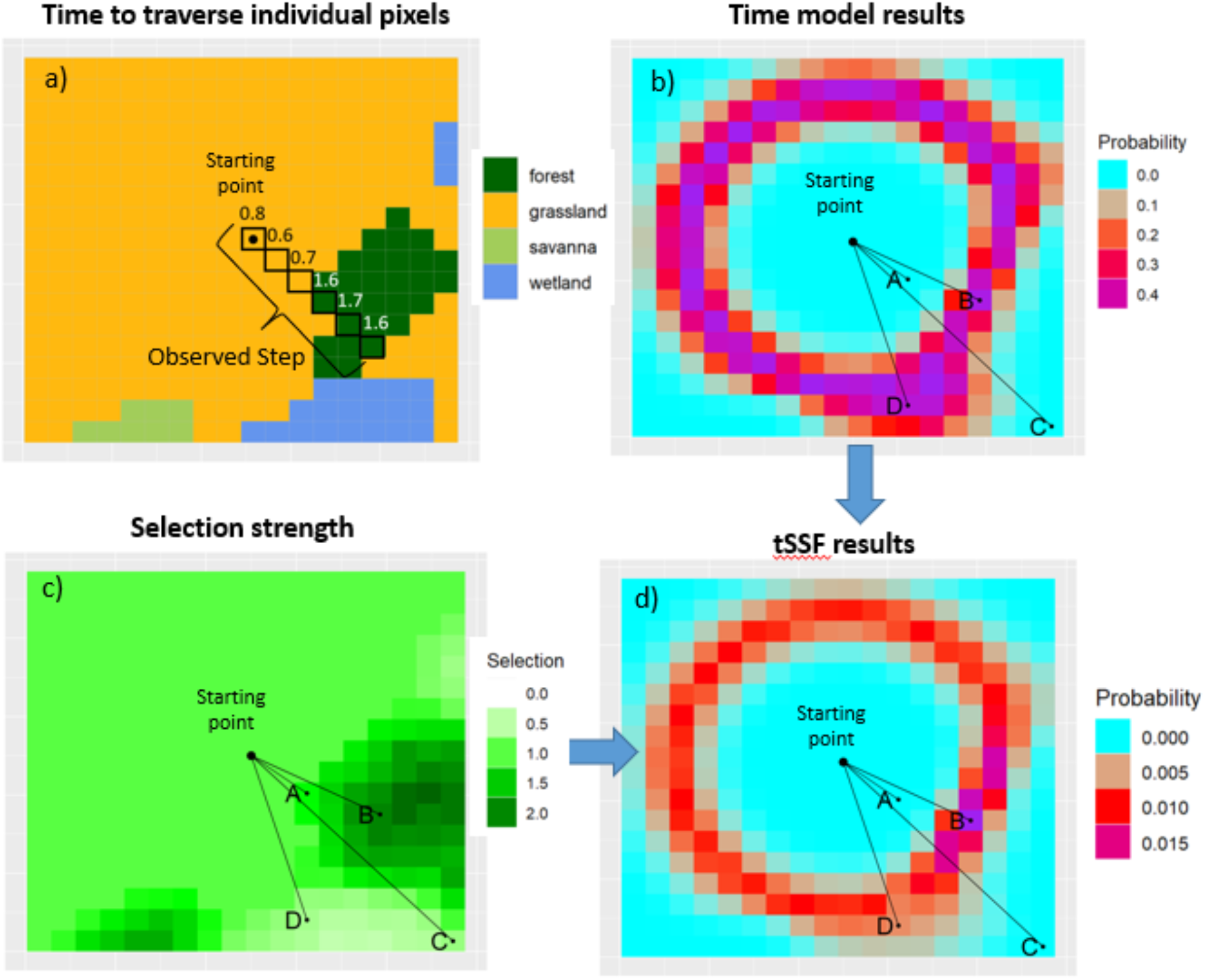
Conceptual description of the time model and the time step selection function (tSSF). (a) A hypothetical landscape illustrating the time taken to traverse a particular path. Traversed pixels are shown with black squares and the time (in minutes) taken to traverse each pixel is shown above the corresponding pixel. (b) Time model results regarding the probability of taking seven minutes to reach each pixel of the landscape given the initial starting point. (c) Selection strength for the different landscape characteristics, where values >1 indicate selection and values <1 indicate avoidance relative to grassland. (d) Selection probabilities based on the tSSF model once the results of the time model and selection strength are taken into account. In panels b, c, and d, we show four potential endpoints (points A - D) in this landscape.

We start by assuming that the time taken to traverse pixel *i* in step *j* (Δ*t*_*ij*_) is given by

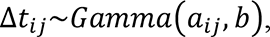

where we assume that 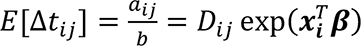. In this expression, *D_ij_* is the distance traveled in pixel *i* in step *j*, 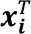 is a vector of covariates associated with pixel *i*, and ***β*** is a vector of regression coefficients. In this gamma regression, time taken to traverse a pixel is assumed to be proportional to the distance traveled but the proportionality constant depends on the characteristics of the pixel. As a result, these slope parameters determine how the mean time taken to traverse a pixel is associated with pixel-level variables such as land-use/land-cover (LULC), elevation, distance to road, or normalized difference vegetation index (NDVI).

Let 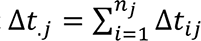 be the total amount of time in step *j*, where *n_j_* is the number of pixels traversed within step *j*. Notice that using standard GPS tracking data, we only observe Δ*t*_.*j*_ while the individual times for each pixel Δ*t*_*ij*_ are latent. For example, in Fig. 2a, the total time taken to traverse these 6 pixels was equal to 7 minutes, but we do not know the time required to cross each individual pixel. Nevertheless, because of the Gamma distribution assumption, it can be shown that

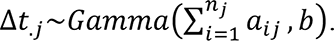

This model can be viewed as representing a gamma process in which, because individual increments arise from gamma distributions, the sum of increments is also gamma distributed. Unfortunately, it can be cumbersome to repeatedly calculate 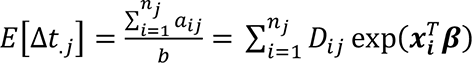 within our model fitting algorithm. To enable this model to be fit in a straightforward fashion, we approximate this quantity by using the mean of the covariates values for the pixels traversed in step 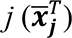 as well as the overall distance in this step (*D*_*j*_):

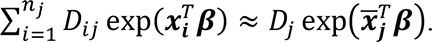

The accuracy of this approximation is likely to be higher if the pixels that characterize the environment are large relative to the step lengths and/or if there is little spatial heterogeneity in the landscape. We test this approximation using simulated data and find that our model works well (see Appendix 1). As a result of this approximation, our model becomes

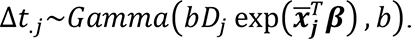

An extension of this basic model allows us to account for additional variability associated with missed fixes. Although tracking devices are often programmed to collect GPS coordinates at regular time intervals, Δ*t*_.*j*_ can be substantially different from the programmed time interval due to missing GPS fixes. When GPS fixes are missed, it is likely that there will be even greater uncertainty regarding the exact path traveled by the animal. For this reason, we modify the above model to allow the variance to potentially increase when GPS fixes are missed

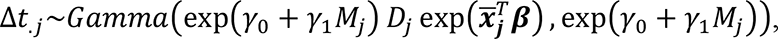

where *M*_*j*_ is a binary variable, equal to 1 if GPS fixes were missing in step *j* and equal to zero otherwise. In this expression, the variance is given by 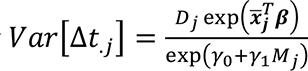. If missed GPS fixes increase the variance, we expect that *γ*_1_ < 0 because the denominator will be smaller when *M*_*j*_=1 and therefore the variance will be larger.

Besides providing inference on how landscape variables (e.g., land use) influence time to traverse a pixel, this model also estimates the probability of reaching different parts of the landscape in a particular time interval, assuming an initial location. For example, Fig. 2b displays the probability that the animal requires seven minutes to reach each pixel in the landscape assuming the animal starts at the center black dot and moves in a straight line. Areas close to the starting point (e.g., point A in Fig. 2b) have low probability because much less than seven minutes are needed to reach these pixels. On the other hand, point C in Fig. 2b also has low probability because much more than seven minutes is required to reach this pixel given that it is very far away from the starting point. Notice that Fig. 2b is asymmetric because, in this example, time taken to traverse the landscape is influenced by the LULC classes along the path. For instance, the probability of taking seven minutes to reach point B is similar to that of reaching point D, despite the fact that point B is closer to the starting point when compared to D. This occurs because the animal moves faster over the path required to reach D (due to the lower proportion of forests and presence of wetlands) whereas the animal moves slower over the path required to reach B (due to the higher proportion of forests and absence of wetlands).

### 2.3. Time-explicit Step Selection Function (tSSF)

To illustrate how the time-explicit step selection function (tSSF) model works, it is useful to refer back to Fig. 2. Figure 2c reveals that the animal might have moved from the starting point to point B but other steps would also have been possible (e.g., the path from the starting point to point D). Furthermore, this figure reveals that, irrespective of its movement capability, this animal tends to select forests and savanna in relation to grassland while avoiding wetlands. Figure 2d shows that the step ultimately chosen by the animal is driven by a combination of the likelihood of the animal reaching that pixel within a given time interval (estimated by the time model) and selection strength for that path irrespective of movement constraints (step selection function).

To more clearly explain the tSSF model, assume that the fix rate from our GPS tracking device is Δ*t*, that the animal is currently in pixel *i* (i.e., *P*_*t*_ = *i*), and that the landscape contains *N* pixels (*i*. *e*., *P*_*t*+Δ*t*_ ∈ {1, …, *N*}). Recall that we are interested in estimating the probability of choosing pixel *j* given time constraint Δ*t* and starting point *i* (i.e., *q*_*ij*_ = *p*(*P*_*t*+Δ*t*_ = *j*|Δ*t*, *P*_*t*_ = *i*)) (Eqn. 1). In this equation, *p*(Δ*t*|*P*_*t*+Δ*t*_ = *j*, *P*_*t*_ = *i*) quantifies the likelihood that Δ*t* will be required to traverse the path between *P*_*t*_ = *i* and *P*_*t*+Δ*t*_ = *j* and can be calculated based on the time model described previously.

We assume that the probability *p*(*P*_*t*+Δ*t*_ = *j*|*P*_*t*_ = *i*) in Eqn. 1 is given by 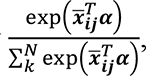 where 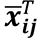 is a vector that contains the mean covariate values in the path from *i* to *j* and ***α*** is a vector containing the corresponding slope parameters. We rely on average covariate values in order to be consistent with the time model formulation, but this is not required (i.e., just the covariate values at pixel *j* could have been used). Our model calculates the probability of moving from pixel *i* to pixel *j* by first specifying the marginal habitat-selection probability and multiplying it by the conditional time probability. This conditional time probability, given by the time model, will automatically distinguish pixels that are available from ones that are not. For instance, a pixel that is too far away for the animal to reach within a particular time interval Δ*t* will be essentially removed from the expression above because *p*(Δ*t*|*P*_*t*+Δ*t*_ = *j*, *P*_*t*_ = *i*) ≈ 0.

It can be computationally intensive to calculate the vector of probabilities for all potential destination pixels because of the large number of pixels across the landscape (i.e., *N* might be very large) (e.g., Aarts et al. 2012, Potts et al. 2014). However, as described in Manly et al. (2002, chapter 8), we can simplify this calculation by selecting the pixel chosen by the animal (denoted as *j\**) as well as randomly selecting *n*-1 pixels not chosen by the animal, resulting in a total of *n* pixels, where *n* is much smaller than *N*. As a result, our likelihood expression becomes

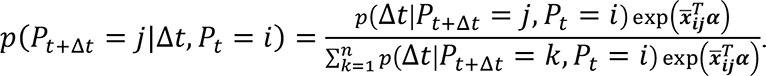

Now the denominator involves a sum over *n* (not *N*) pixels, where *n* ≪ *N*. This is the expression of a conditional logistic regression that explicitly accounts for time. Notice that the set of available steps can be sampled in various ways (i.e., it is not necessary to rely on samples from the empirical distribution of step lengths). This is supported by statistical theory (see chapter 8 in Manly et al. 2002) and is confirmed by our simulations (see Appendix 2). Ultimately, the data to be used for the tSSF model will be comprised of the selected steps with the corresponding covariate information, matched with one or more available steps and their associated covariate variables. Furthermore, the tSSF model will also require the corresponding pre-computed time model probability for each step, regardless if it is an available or selected step.

### 2.4. Empirical analysis: Giant Anteater case study

#### 2.4.1. Movement data

The data were collected between 2013 and 2017 in a 350-km^2^ area in the Brazilian Pantanal (19°16′60ʺS, 55°42′60ʺW). The landscape is a mosaic of habitats that include forests, open grassland, pasture, savannas, and wetlands (Fig. 3). Historical mean temperature is 25.4°C and climate is classified as semi-humid tropical (“Aw” in Köppen’s climate classification). Traditional extensive cattle ranching is practiced in the area, but overall anthropogenic impacts and threats are relatively low.

**Fig. 3.**
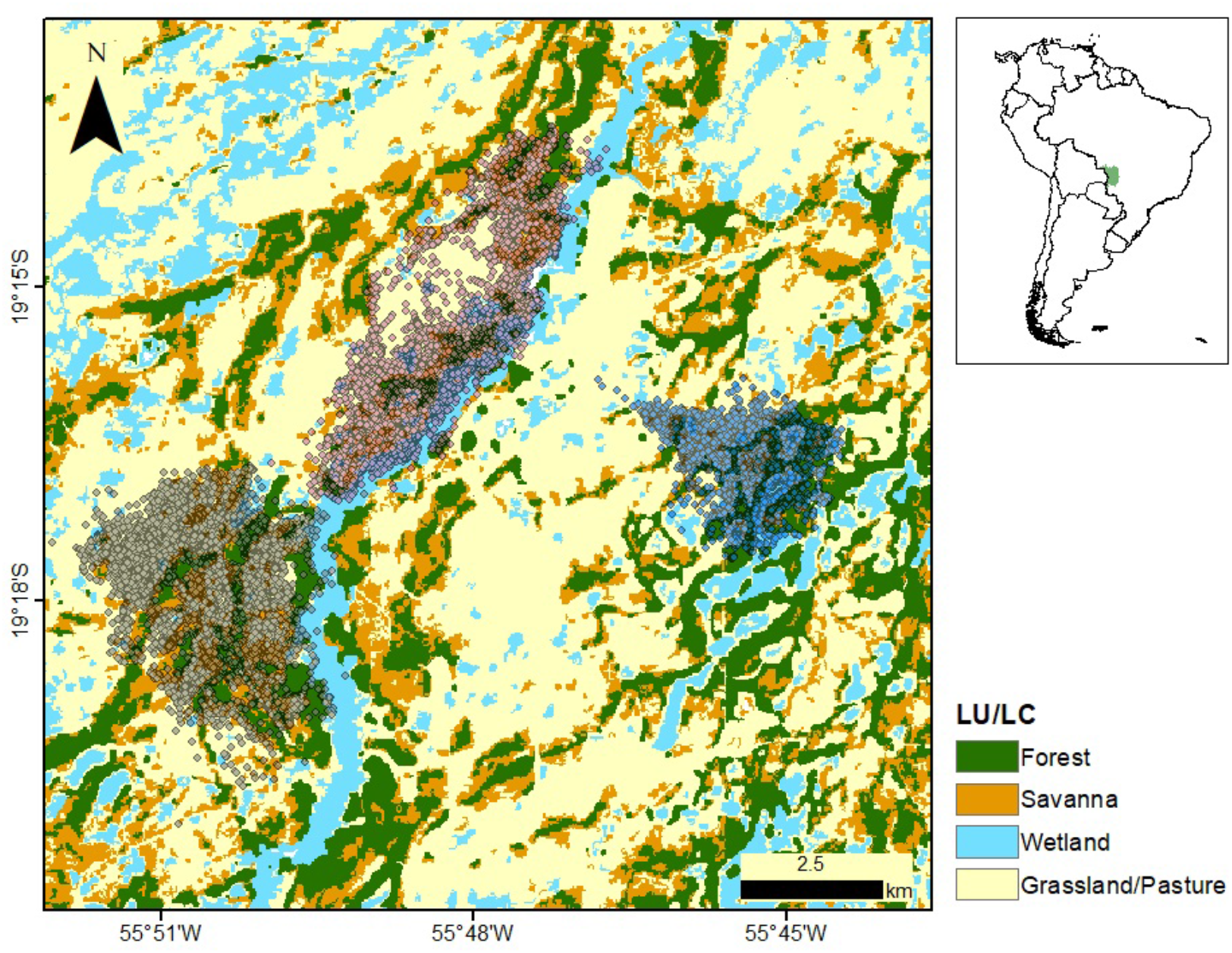
Land-use land-cover (LULC) classification of the study region for the year 2016 according to Mapbiomas (https://mapbiomas.org/en). Three individual giant anteaters (*Myrmecophaga tridactyla*) were monitored through GPS telemetry (semi-transparent circles) in the Brazilian Pantanal wetlands (green polygon on the inset).

Anteaters were captured, immobilized, and sedated following the procedures described in Kluyber et al. (2021). Each individual was sexed, weighed, and equipped with a global positioning system (GPS; TGW-4570, Telonics) harness. None of the tracking devices exceeded 3% of the animals’ body mass. The GPS devices were programmed to record location points at 20 or 30-min intervals (depending on the animal). However, because some GPS fixes failed to be acquired, the time interval between fixes was sometimes greater than 20 or 30 minutes. As the time interval increases (i.e., as the number of failed GPS fixes increase), the assumption of a straight-line path becomes less reliable. For this reason, we removed observations for which the time interval was greater than one hour and, whenever the time interval was greater than 35 minutes (to allow for some tolerance around the 30 min time interval), we allowed for increased variance in our time model by setting *M*_*j*_ to one.

We also removed observations with speeds unlikely to be achieved by the species (> 0.33 m/s, approx. the 99^th^ percentile of speed). Taken together, the removal of observations with very large time intervals or with unrealistic speeds resulted in the elimination of 2% of the data. Also, an important observation is that our time model is not defined when distances are exactly equal to zero. Therefore, whenever the distance between two GPS fixes was equal to zero (0.2% of our observations), we set distance to the smallest non-zero distance that was recorded (i.e., 1 meter). The final movement dataset contained ∼65,000 observations from three individuals (Table 1).

**Table 1.** Summary of the movement data used by the time and the tSSF models. F and M stand for female and male, respectively.

| ID | Sex | Monitoring period |  | Duration<br>(days) | Number of observations |  |
| --- | --- | --- | --- | --- | --- | --- |
|  |  | Start | End |  | Time model | tSSF |
| Berenice | F | Oct 2015 | Nov 2016 | 386 | 26,996 | 15,577 |
| Brigite | F | Jul 2013 | Jul 2014 | 340 | 11,233 | 7,510 |
| Fergus | M | Jul 2016 | Aug 2017 | 407 | 27,010 | 13,115 |
| **Total** |  |  |  | **1,133** | **65,239** | **36,202** |
| **Average** |  |  |  | **378** | **21,746** | **12,067** |

To characterize the habitat, we relied on Collection 5 of land-use land-cover classification (LULC) provided by Mapbiomas for the Pantanal ecosystem (https://mapbiomas.org/en) for each year between 2013 to 2017. Pixel size of this LULC product is 30 x 30 m. We also relied on hourly temperature data collected by the nearest automatic meteorological station of the National Institute of Meteorology of Brazil (INMET). Given that these temperature data occasionally exhibited abnormal temporal patterns (e.g., sudden drops followed by sudden increases of temperature) and there were some missing data, we decided to rely on the median temperature for each hour in each month and each year as a more robust measure of temperature that reflects both within day variation as well as seasonal variation.

#### 2.4.2. Fitting the time model

For each step, defined as the straight line between two consecutive locations, we extracted the proportion of each LULC cover within a 30 m buffer of the path to account for GPS location and individual path uncertainty (Zeller et al. 2016). We combined the grasslands and pasture classes (hereafter grasslands), and then used it as the baseline LULC. As a result, we only include the proportion of forest, savanna, and wetland as covariates in our time model. Finally, to account for diel patterns in movement, we relied on cyclic cubic B splines to characterize how time taken to traverse a particular path depends on time of day, where the knots were set to 6 am, 12 pm, and 6 pm.

We fit this model in a Bayesian framework using JAGS (Plummer 2003). Separate models were fit for each individual. Vague priors were adopted for *γ*_0_, *γ*_1_, and *β*_0_ whereas we used more conservative priors for the slope parameters *β*_*p*_ (p=1,…,P):

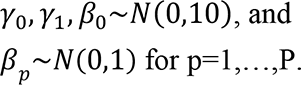

A tutorial describing how to prepare the data and fit the time model is provided in Appendix 3.

#### 2.4.3. Fitting the tSSF model

To create the available steps that the animal could have taken, we defined four available steps of the same length as the observed step, one for each cardinal direction (i.e., east, west, north, and south from where the step started). Similar to the analysis for the time model, the proportion of each LULC class in the area surrounding each step was calculated by creating a 30-m buffer around the straight line that connects two consecutive locations and determining the proportion of pixels associated with each LULC class. Furthermore, although grasslands/pastures are present in the study region, we did not include this LULC class as covariate in the model because they act as the baseline LULC class. Finally, we removed observations with missing temperature data and steps for which LULC was identical for the observed and available steps (sample size for each individual is given in Table 1). The reason for this last procedure is that there is no information on selection strength if the habitat information is the same for the observed and available steps because these cancel out when calculating the SSF ratio (McDonald et al. 2006, Vardakis et al. 2015).

We also fit this model in a Bayesian framework using JAGS (Plummer 2003). Separate models were fit for each individual using the following conservative priors for the slope parameters:

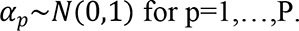

A tutorial describing how to incorporate the time model results and fit the tSSF model is provided together with this article (Appendix 3).

#### 2.4.4. Connectivity implications

We also examine the implications of the tSSF results in terms of characterizing permeability in heterogeneous and fragmented landscapes. To estimate the amount of flow of individuals through heterogeneous landscapes, the expected number of individuals in each pixel after *t* time steps was calculated as N**Ψ**^T^**Q**^*t*^, where *N***Ψ** characterizes the initial distribution of individuals in each pixel of the landscape (Fletcher et al. 2019). To illustrate these time-explicit calculations, we created a hypothetical landscape composed by a large wetland surrounded by grasslands with two patches of savanna and estimated the flow of individuals at different points in time. We assume that 100 individuals start at one savanna patch at the beginning of the simulation aiming to reach another savanna patch. To set a savanna patch to be the destination, we selected a pixel *i*^∗^ at the center of this patch and we modified the **Q** matrix by setting *q*_*i*∗*j*_ = 0 for *i*^∗^ ≠ *j* and *q*_*i*∗*i*∗_ = 1.

## 3. Results

### 3.1. Time model results

The results for the time model applied to the simulated data showed that our model was able to estimate the true parameters well despite relying on an approximation where covariates are averaged along each step, even for landscapes that are more spatially heterogeneous (Appendix 1). The results for the time model applied to the giant anteater data show that individuals consistently moved slower when traversing forests and savannas and faster when traversing wetlands, in comparison to the time taken to traverse grasslands (the baseline LULC) (Fig. 4a). Furthermore, based on our cyclic splines, we also find that giant anteaters tended to move much slower during daytime, from approximately 5 am to 8 pm, indicating that this species tends to have a nocturnal/crepuscular activity pattern (Fig. 4b). Finally, as expected, the *γ*_1_ coefficients associated with the missed GPS fixes were consistently estimated to be negative, revealing that missed GPS fixes resulted in greater uncertainty in our time model (see Appendix 4).

**Figure 4.**
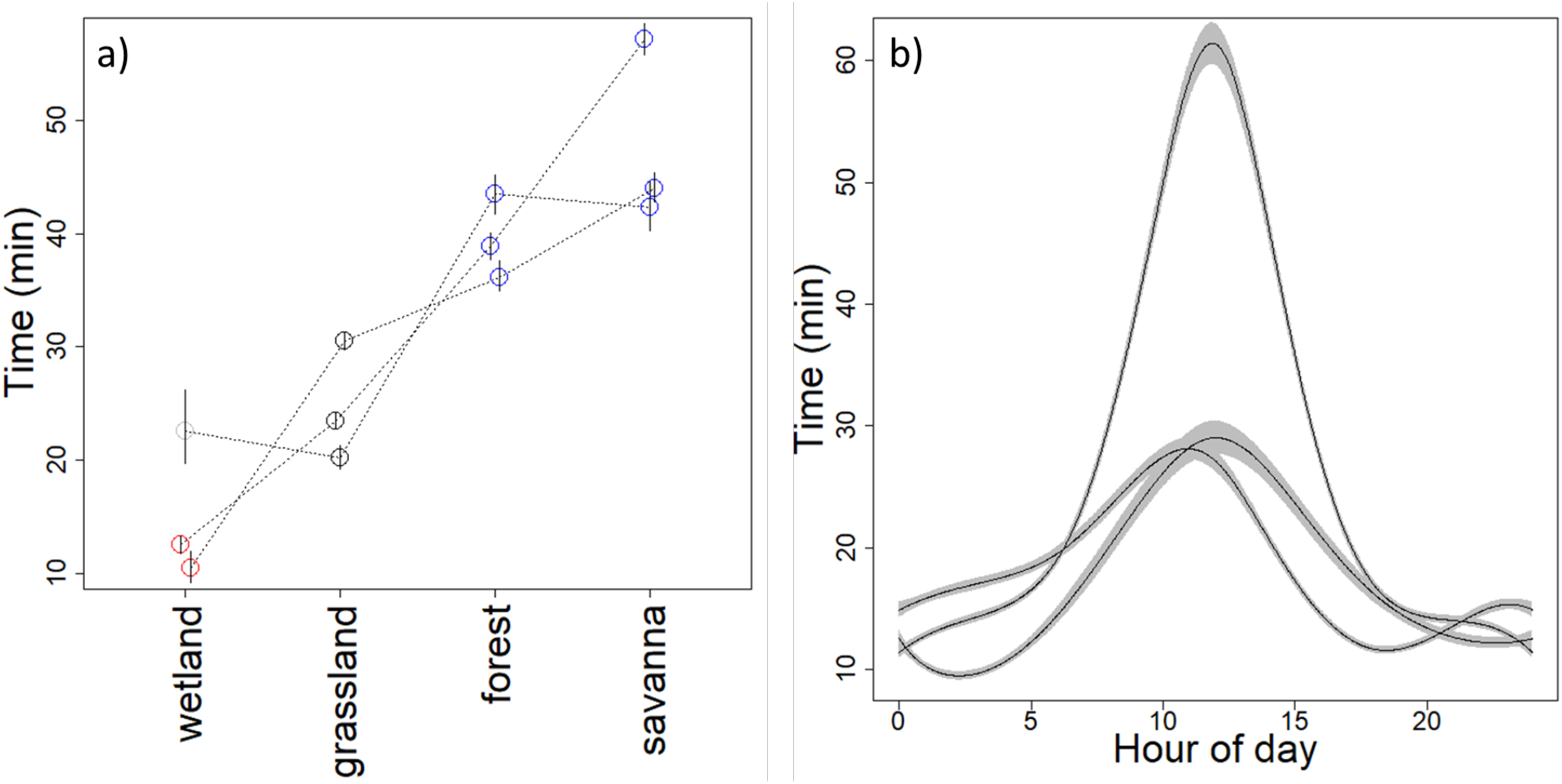
Results from the time model. (a) Estimated mean time (in minutes) taken to traverse 50 m in different types of habitat at 8 pm. Each circle represents the result for a given animal and LULC category. Circles connected by the same line correspond to posterior median results from the same individual. Blue and red circles denote statistically positive (i.e., p(*β*<0)<0.025) and negative effects (i.e., p(*β*>0)<0.025), respectively, whereas grey circles show results that are not statistically discernible from grasslands (the baseline LULC class). Vertical lines are 95% credible intervals. (b) Estimated mean time required to traverse 50 m in the baseline LULC class (grasslands) throughout the day, showing that animals moved slower and were more likely to be inactive during the daytime. Each line corresponds to the posterior median for an individual animal and gray ribbons are the corresponding 95% pointwise credible intervals.

### 3.2. Time-explicit step selection function (tSSF) results

The results for the tSSF model applied to the simulated data reveals that this model can estimate the true parameters well, regardless of the number of available steps and the method used to choose these steps (Appendix 2). The tSSF model applied to the giant anteater data reveals that wetlands are avoided by all individuals relative to grasslands (the baseline LULC class) at the mean temperature of 25 °C (Fig. 5a). Although the selection for forests and savanna is ambiguous relative to grasslands, there is a consistent pattern of increased selection strength from wetlands to forests to savannas at mean temperature. Interestingly, the parameter estimates for the interaction between forest and temperature were consistently positive (Fig. 5b), indicating that selection strength for forests in relation to grasslands tends to increase with increasing temperatures. A similar pattern seems to hold for savanna, but generally the interaction is less strong and the result for one of the individuals was not statistically discernible from zero.

**Figure 5.**
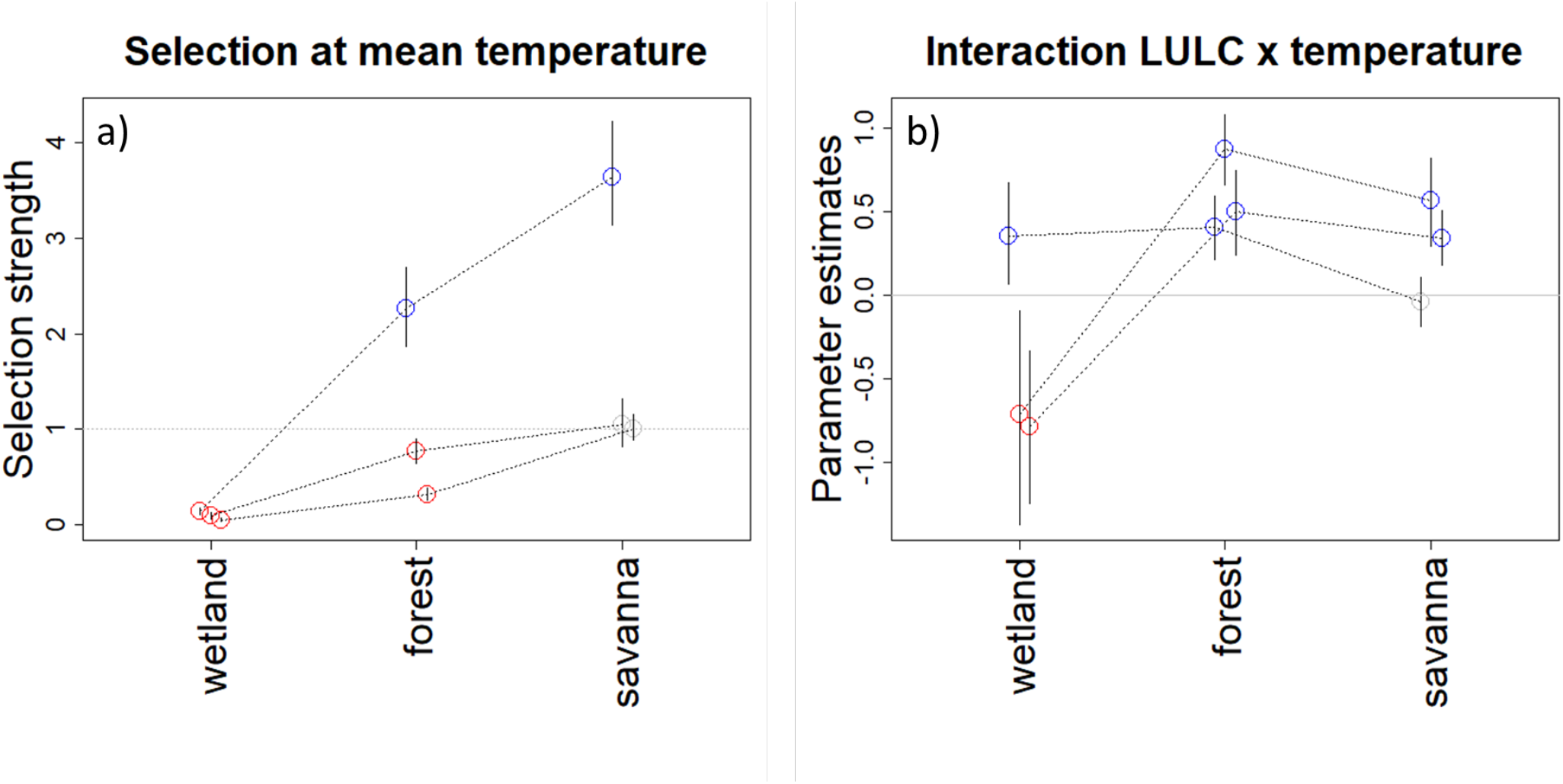
Estimates of (a) selection strength for different types of habitat at mean temperature (calculated as 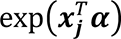) and (b) the effect of the interaction between LULC and temperature on selection strength. The horizontal grey line depicts the results for grasslands (the baseline LULC category) in panel A. Each circle represents the posterior median result for a given animal and LULC category and lines connect results from the same animal. Blue and red circles denote statistically positive (i.e., p(*α*>0)>0.975) and negative effects (i.e., p(*α*<0)>0.975), respectively, whereas grey circles indicate estimates that are not statistically discernible from zero. Vertical lines are 95% credible intervals.

Selection strength (calculated as 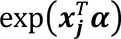) can also be displayed spatially given that it depends on LULC (top panels in Fig. 6), clearly revealing how individuals consistently avoid wetlands and how selection strength for different landscape features dramatically changes as temperature increases (middle and bottom panels of Fig. 6)

**Fig. 6.**
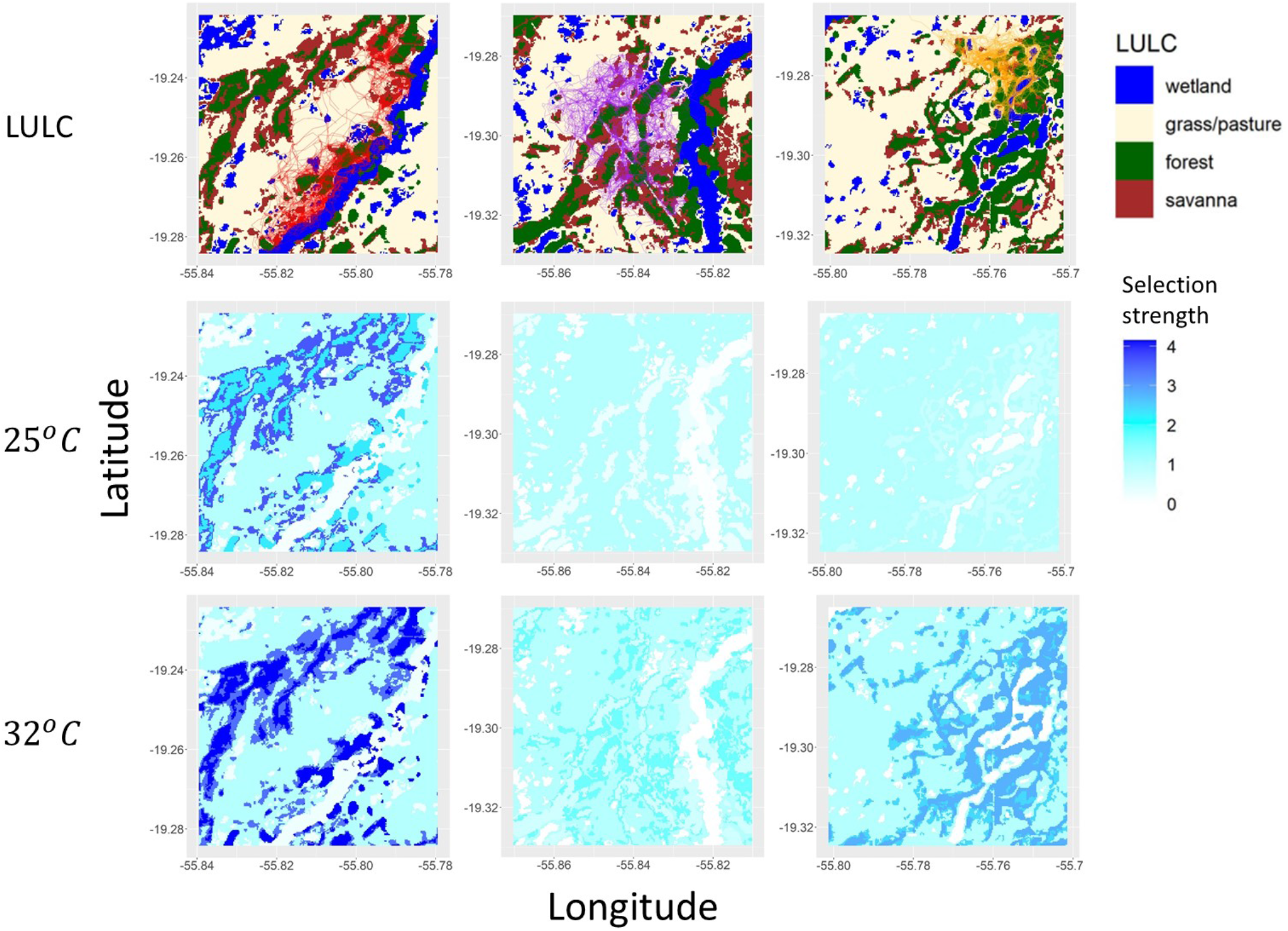
Influence of temperature in the selection strength of landscape features. Top panels show the spatial distribution of LULC together with the corresponding animal tracks. Middle panels show selection strength across the landscape based on mean temperature (25°C) whereas bottom panels show selection at a temperature that is 1.5 standard deviation above the mean (32°C). In this figure, selection strength is calculated as 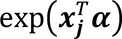 and selection for grasslands is equal to one because this is the baseline LULC. Each column of panels presents the results for one of the individual anteaters (*Myrmecophaga tridactyla*) that were tracked.

Combining the time model with the tSSF results enabled the characterization of landscape permeability and selection strength under different temperature scenarios. Figure 7 reveals that giant anteaters consistently move faster and avoid wetlands relative to grasslands, regardless of temperature (lower left quadrant). Furthermore, the individuals in our dataset generally selected for forests and savannas relative to grasslands, particularly at higher temperatures (upper right quadrant). These results suggest that animals rely on forests and savannas for slower behaviors (e.g., resting or foraging) and are more likely to increase their selection for these habitats as temperatures rise.

**Fig. 7.**
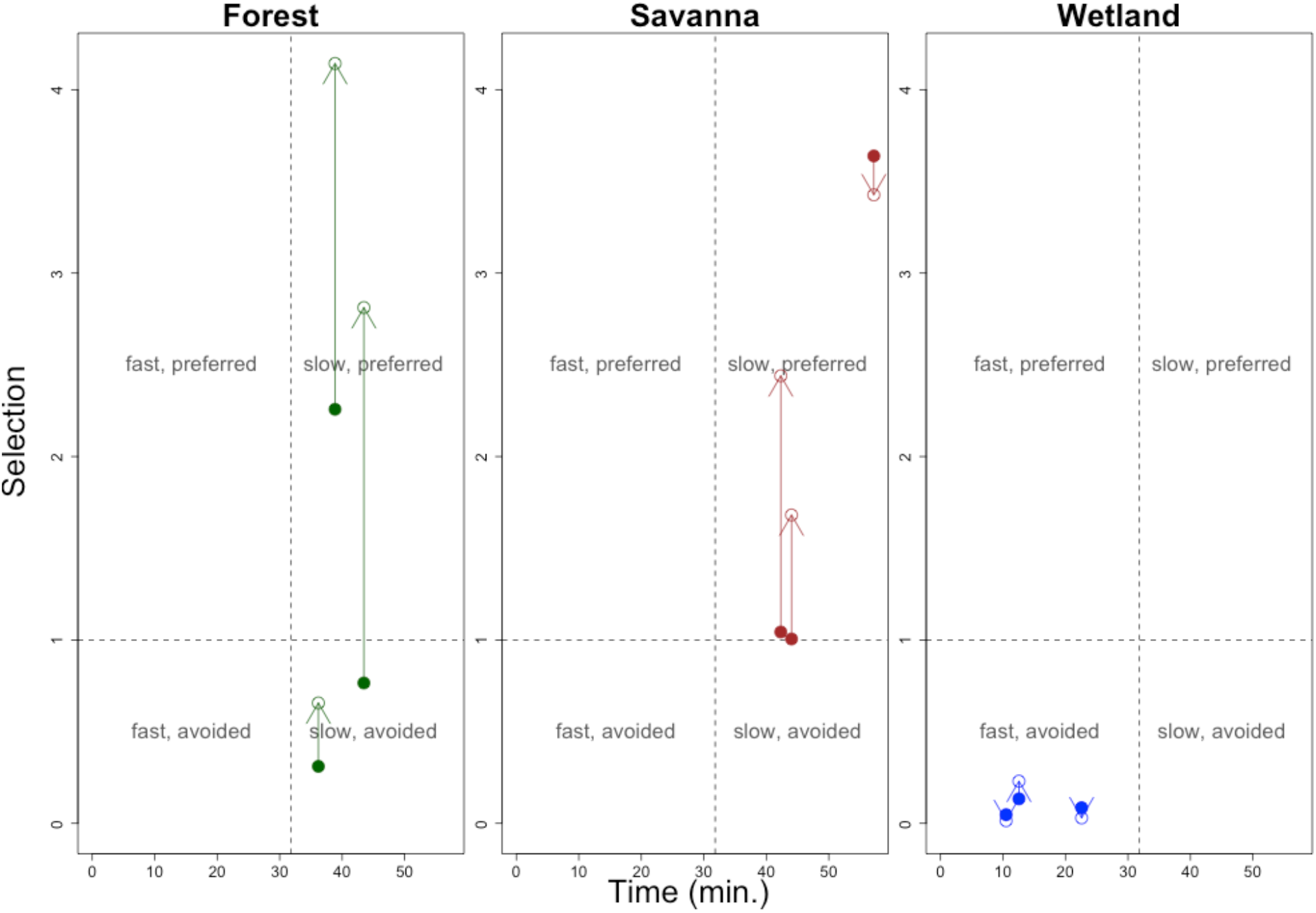
Characterization of landscape use by giant anteaters relative to time and selection strength (calculated as 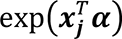) at two temperature scenarios. Each point represents the result for a particular individual in a given temperature scenario, and each panel shows the results for a LULC category. Estimated mean time value for each individual refers to the time taken to traverse 50 m of the LULC category at 8 pm. Arrows indicate how selection strength is likely to change with increasing temperatures, connecting selection strength values at the mean temperature (25 ℃; solid circles) and at 1.5 standard deviation above the mean (32 ℃; open circles). The dashed vertical line represents the mean of the time estimates to traverse all LULC categories, whereas the dashed horizontal line represents the selection strength for grasslands (the baseline LULC category).

### 3.3. Implications for connectivity

Recall that the simulated landscape (Fig. 8) contains a large wetland (ellipse) surrounded by grasslands with two patches of savanna (rectangles) and that individuals start at the upper left patch (origin patch) and end up in the patch to the right of the wetland (destination patch). Our time-explicit predictions reveal that individuals do not use the shortest path to the destination patch because that would have required them to traverse the wetland, an avoided habitat type. Instead, these individuals tend to move around the wetland to eventually reach the destination patch (Fig. 8a). Importantly, our results suggest that approximately 49 days are required for 90% of the individuals to reach the destination patch (Fig. 8b).

**Fig. 8.**
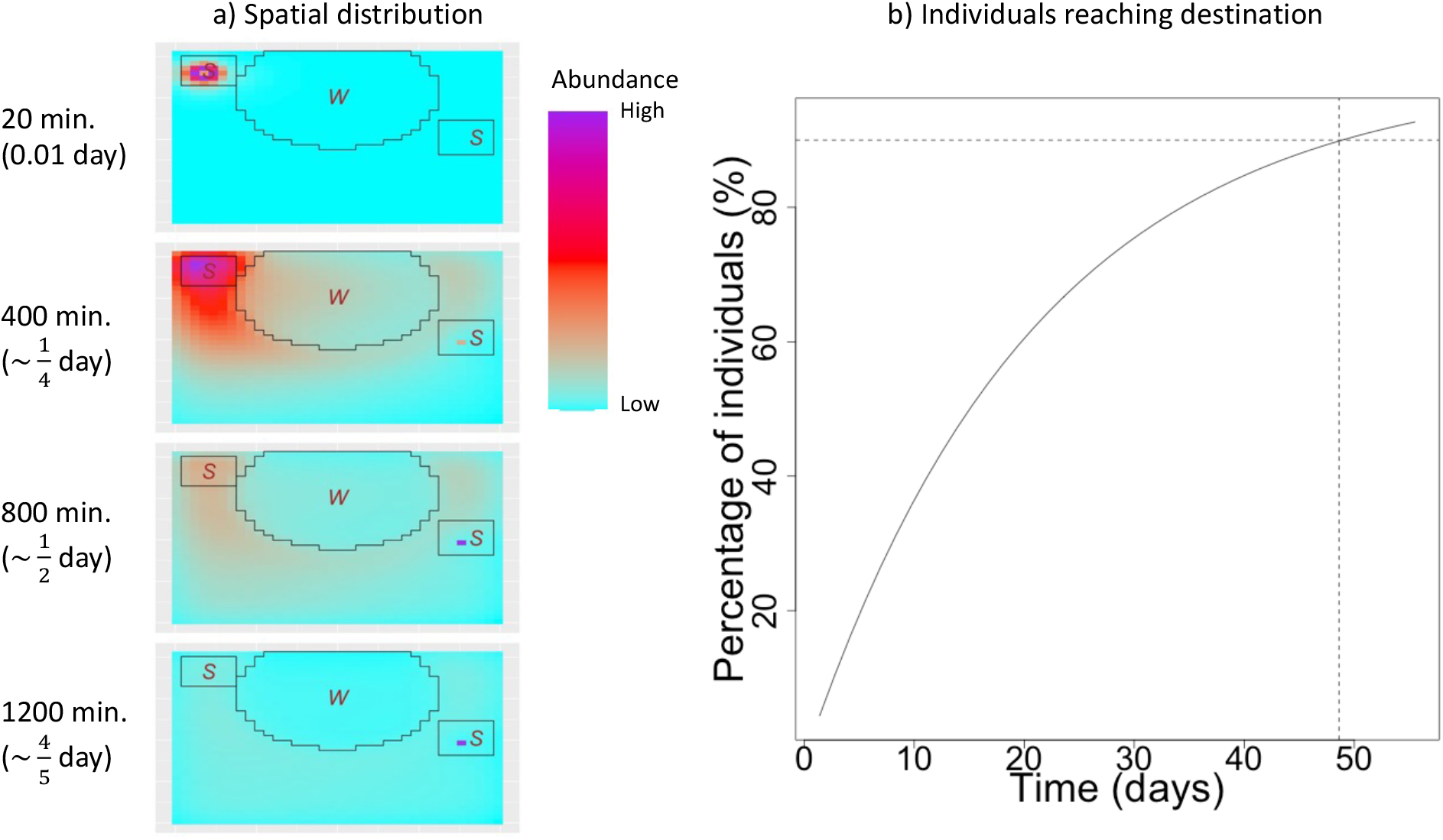
Connectivity implications of the inferred time and selection processes for a simulated landscape. This landscape consists of a wetland (ellipse with “W”) and patches of savanna (rectangles with “S”) within a grassland matrix (rest of the area). One hundred individuals start in the upper left savanna patch at time 0 and each panel in (a) shows the predicted abundance of individuals on each pixel after 1, 20, 40, and 60 time steps, where each time step corresponds to 20 minutes. The color gradient indicates predicted abundance of individuals, but note that the scale is not the same across different panels. Results in (b) show the estimated percentage of individuals in the destination patch as a function of time. Horizontal and vertical dashed lines highlight that approximately 49 days are required for 90% of the individuals to have reached the destination patch. All results are based on the estimated parameters for a single individual, assuming movement patterns at 8 pm and at mean temperature (25℃).

## 4. Discussion

In this article, we have proposed two novel models, the time model and the time-explicit step selection function (tSSF), and we have shown how these models can provide complementary information by explicitly distinguishing the drivers of time from the drivers of selection in animal movement. Furthermore, we have shown how tSSF can be integrated into frameworks focused on estimating landscape connectivity without requiring the arbitrary transformations that plague connectivity analysis, resulting in time-explicit predictions. Below we compare the tSSF to previous habitat selection models and discuss our findings based on the giant anteater data from the Pantanal region.

### 4.1. Comparison with previous modeling approaches

The time model proposed in this study is different from earlier approaches because it focuses on modeling time rather than distance, speed, or velocity. Our focus on modeling time is an important characteristic for two reasons. First, many connectivity problems inherently require time-explicit solutions (e.g., if species can track a changing climate) and linking movement model results to a time-explicit framework (e.g., SAMC) requires a model for time (not distance, speed, or velocity) to be able to calculate the elements *q*_*ij*_ = *p*(*P*_*t*+Δ*t*_ = *j*|Δ*t*, *P*_*t*_ = *i*) in the permeability matrix **Q**. Second, assuming a gamma process and a straight-line assumption, this model enables inference of the time taken to traverse individual pixels and the factors that influence this time. On the other hand, speed (or velocity) at the step scale is not easily translated into speed (or velocity) at the pixel level, precluding the understanding of how speed (or velocity) relates to pixel-level characteristics.

The tSSF model contains a landscape-dependent kernel (i.e., the time model), which describes the probability of requiring a given amount of time to cross a heterogeneous landscape. As a result, this kernel can capture the fact that some landscape characteristics may favor faster movements while other characteristics might favor slower movements. In contrast, standard SSF models (Forester et al. 2009) as well as models that attempt to link resource selection and step selection models (Michelot et al. 2019) rely on resource-independent movement kernels. While some authors have suggested the inclusion of an interaction between step length and landscape characteristics in RSF and SSF to account for spatial heterogeneity in movement rates (e.g., to account for higher movement rates in open habitats compared to forested areas; Nielson et al. 2009), this is seldom done and conflates inference on the drivers of selection strength with inference on the drivers of speed.

The tSSF model is most similar to the integrated step selection analysis (iSSA) described in Avgar et al. (2016) but there are important differences. First, iSSA can generate parameter estimates for the movement kernel that are nonsensical. For example, the shape and scale parameters of the Gamma distribution (assuming this is the distribution chosen for modeling step length) cannot be negative. However, because these parameters are modeled as linear functions of covariates within the current formulation of iSSA, the resulting shape and scale parameters can actually be estimated to be negative even if the data generating mechanism relied on positive parameter values (see simulation results in Appendix 5). Second, parameter estimability is an important problem in iSSA and movement parameters exhibit compromised precision even when using relatively large data sets (Avgar et al. 2016) (see simulation results in Appendix 6). Our model formulation and two-stage model fitting approach are able to avoid both of these problems. By relying on Bayes theorem, the tSSF is a much more principled approach to quantifying the probability of moving from one pixel to another than iSSA.

### 4.2 Giant anteater case study and connectivity implications

The time model applied to data from giant anteaters in the Pantanal region revealed that individuals tended to be most active between 8 pm and 5 am. Nocturnal behavior has been recorded for this species, especially on warmer days (Camilo-Alves and Mourao 2005, Mourao and Medri 2007). Furthermore, we found that individuals tended to consistently move faster over wetlands, possibly because these environments are relatively poor in feeding resources when flooded and provide little vegetation cover, which increases predation risk. This hypothesis seems to be corroborated by the tSSF results, which revealed that wetlands were consistently avoided relative to all other land cover classes. In contrast, giant anteaters tend to move slowest over forests and savannas. This slower movement might be associated with not only increased biomechanical resistance offered by more vegetation, but also the fact that these habitats are used for foraging and resting (Bertassoni and Ribeiro 2019, Giroux et al. 2021b). Indeed, the tSSF model showed increased selection strength for forests and savannas, particularly at higher temperatures. Previous studies suggest that forests may act as a thermal shelter for giant anteaters, not only when temperatures are high and above their thermoneutral zone, but also when temperatures are very low (Camilo-Alves and Mourao 2005, Giroux et al. 2021a). One of the challenges of determining the effect of temperature on animal behavior is that it is highly correlated with the time of the day. In this study, we chose to incorporate hour of day in the time model, whereas temperature was included in the step selection function. However, a more complete understanding of the effect of temperature on individual behavior will require additional studies to better disentangle these processes.

By interpreting time and selection strength as distinct axes that govern animal behavior, we were able to characterize each LULC class in a biologically meaningful way (see Box 1) that can have important implications for conservation. For example, wetlands consistently fell in the “fast, avoided” quadrant, regardless of temperature. In contrast, we find that forests and savannas tend to fall in the “slow, selected” quadrant, particularly with higher temperatures, suggesting that, while these habitats might not favor fast movement, they may be critical for the foraging and resting of giant anteaters, being good to compose slow corridors (Bastille-Rousseau and Wittemyer 2021). Interestingly, none of our estimates fell in the “fast, selected” quadrant (upper left quadrant), which could potentially be the prime target when designing wildlife corridors (LaPoint et al. 2013). We hypothesize that this might be due to the fact that all of the individuals in our dataset were resident individuals and dispersing individuals might have important differences in terms of speed and habitat selection when compared to resident individuals (Beier et al. 2008).

Part of our contribution is to show that the tSSF model results can be used to directly populate models for connectivity assessments rather than using resistance maps that require arbitrary decisions to relate measures of habitat quality (e.g., resource selection strength) to landscape resistance. Critically, we show that using the tSSF model together with SAMC allows us to generate time-explicit predictions of dispersal patterns, something that many connectivity models cannot represent. We note, however, that our dispersal example has several limitations. First, this example is based on just one individual, not all three individuals in our dataset. Second, our results are based on tSSF estimates for a particular time of day and temperature. Similar to many connectivity analyses (Kumar and Cushman 2022), our results do not account for individual or temporal variability, such as diel patterns and seasonal changes in activity level and temperature. Third, we do not take mortality risk into account even though this is a critical piece of information when understanding landscape use (Fletcher et al. 2019). Finally, it can be computationally challenging to scale up our connectivity analysis because we allow for transitions beyond the 8 nearest neighbors, resulting in a much denser (i.e., less sparse) matrix than most connectivity applications (Fletcher and Fortin 2018).

### 4.3. Model limitations and future improvements

Our proposed models are better suited for shorter time-steps because we assume a linear path between GPS fixes. This linear path assumption is arguably the strongest assumption that our models make, and this assumption directly influences a) the landscape covariate values that are used within the model; and b) our estimate of the distance traversed. To take into account the fact that the exact path between two GPS fixes is unknown and therefore the environmental characteristics may not correspond to those in a straight line, we have used a 30-m buffer around this straight line. One could also potentially use Brownian motion/diffusion to better characterize the environment (Horne et al. 2007). This would be particularly useful when one or more GPS fixes are missing because, in these cases, a much larger area would have to be considered to properly characterize the environment. Alternatively, a potential path taken by the animal could be sampled after fitting a continuous-time correlated random walk through the “crawl” R package (Johnson et al. 2008). Finally, a third option would be to restrict the analysis of the time model to steps that occur in relatively homogeneous landscapes. In this situation, there is much less ambiguity regarding the characteristics of the environment that was traversed.

In relation to traversed distance, the assumption of a straight-line path almost certainly leads to an underestimate of this distance, but there are relatively limited options to circumvent this problem at the moment. Aside from sampling a potential path using the crawl R package (Johnson et al. 2008), another option would be to use a dead-reckoning approach to better approximate the actual path taken by the individual (Bidder et al. 2015, Munden et al. 2021). Unfortunately, this approach relies on specialized sensors (e.g., accelerometers and magnetometers) that are still relatively uncommon in the field, requires estimation of speed (e.g., as a function of dynamic body acceleration [DBA]), and the required calculations can be challenging to implement (but see Gunner et al. 2021). Nevertheless, if and when dead-reckoning becomes a more commonly adopted method, the time model will still be useful and will yield more realistic results by not having to rely on a straight path assumption. For the moment being, as long as step length is still a good proxy for distance traveled (as assumed by several models commonly used in movement ecology such as hidden Markov models that rely on step lengths (Langrock et al. 2012, Patterson et al. 2017, Pohle et al. 2017) and iSSA (Avgar et al. 2016)), the results of the time model are likely to be insightful and useful.

Another important limitation is that, similar to the distributions that are often used to model step length (e.g., gamma distribution) in Hidden Markov Models and iSSA (Avgar et al. 2016), the time model assigns zero probability for distances that are equal to zero. This is not a problem from the perspective of model fitting because very few observations had distances exactly equal to zero. Indeed, even if the monitored individual is not moving, the distance between two GPS fixes is typically positive because of geolocation error. However, this characteristic can be problematic when inferring dispersal and connectivity patterns because it assumes that the monitored individual never stays in the same location for two consecutive time-steps. To circumvent this issue, the time model could be modified to include a two-stage process. In the first stage, the animal decides to either stay in the same location or move. If the animal decides to move, then the time model can be used to understand the time needed to traverse landscapes with different characteristics. Such a model would require distinguishing from the GPS tracking data when the animal is not moving from when the animal is exhibiting limited movement. Extending the time model to account for this two-stage process is an important area for future research.

## 5. Conclusions

Landscapes across the world are changing at ever increasing rates due to habitat degradation and loss associated with land use change. Connectivity analysis plays a central role in mitigating these impacts by identifying potential dispersal routes and corridors but better connecting these analyses with movement data is paramount to ensure the reliability of its results. Furthermore, time-explicit predictions of the flow of individuals through the landscape are critical to many connectivity problems but few modeling frameworks can generate such predictions. The methods proposed here can advance the understanding of the ecological and functional roles of different habitat features for species, deepening our understanding of animal habitat selection patterns, and can improve how movement-based modeling results are incorporated into connectivity analysis, resulting in time-explicit predictions.

## 6. List of abbreviations

LULC: Land-Use/Land-Cover
tSSF: Time-explicit Step Selection Function
iSSA: Integrate Step Selection Analysis
GPS: Global Positioning System
SAMC: Spatial Absorbing Markov Chain (SAMC)
JAGS: Just Another Gibbs Sampler
NDVI: Normalized Difference Vegetation Index
INMET: National Institute of Meteorology of Brazil
DBA: Dynamic Body Acceleration

## 7. Declarations

### Ethics statement

All animal handling and monitoring procedures were conducted in accordance with the Guidelines of the American Society of Mammalogists for the use of wild mammals in research (Sikes et al., 2016) and were performed under the license number SISBIO 38218–1 (Chico Mendes Institute for Biodiversity Conservation).

### Consent for publication

Not applicable.

### Availability of data and materials

Movement data for the giant anteaters are available for visualization in Movebank (www.movebank.org with study name “Myrmecophaga tridactyla Pantanal”). Data download requests may be sent to the Center for Species Survival Brazil (SSC) from the International Union for Conservation of Nature (IUCN). The LULC class data are freely available at https://mapbiomas.org/en/download whereas the temperature data are available from the Brazilian National Institute of Meteorology (INMET; https://portal.inmet.gov.br/dadoshistoricos).

### Authors’ contributions

D.V. and N.A. jointly conceptualized the article, created figures, and wrote the original draft. D.V. developed the model with input from J. C. and M. H. and ran the analysis. A.G. and L.G.R.O.-S. provided important suggestions and feedback regarding the giant-anteater case study. A.L.J.D. acquired funding for, participated on, and supervised the giant-anteater data collection. R.F. provided critical suggestions and feedback regarding movement and landscape ecology, particularly regarding the spatial Markov chain framework. All authors reviewed and edited the manuscript, substantially improving it.

## Acknowledgements

We thank G. Massocato and D. Kluyber for helping to capture and monitor the giant anteaters and the Baia das Pedras. We also thank neighboring ranches for allowing us to conduct this study on their land.

## References

Aarts, G., J. Fieberg, and J. Matthiopoulos. 2012. Comparative interpretation of count, presence-absence and point methods for species distribution models. Methods in Ecology and Evolution 3:177–187.

Abrahms, B., S. C. Sawyer, N. R. Jordan, J. W. McNutt, A. M. Wilson, and J. S. Brashares. 2016. Does wildlife resource selection accurately inform corridor conservation? Journal of Applied Ecology 54:412–422.

Avgar, T., A. Mosser, G. S. Brown, and J. M. Fryxell. 2013. Environmental and individual drivers of animal movement patterns across a wide geographical gradient. Journal of Animal Ecology 82:96–106.

Avgar, T., J. R. Potts, M. A. Lewis, M. S. Boyce, and L. Börger. 2016. Integrated step selection analysis: bridging the gap between resource selection and animal movement. Methods in Ecology and Evolution 7:619–630.

Bastille-Rousseau, G., and G. Wittemyer. 2021. Characterizing the landscape of movement to identify critical wildlife habitat and corridors. Conservation Biology 35:346–359.

Beier, P., D. R. Majka, and W. D. Spencer. 2008. Forks in the Road: Choices in Procedures for Designing Wildland Linkages.. Conservation Biology 22:836–851.

Bertassoni, A., and M. C. Ribeiro. 2019. Space use by the giant anteater (Myrmecophaga tridactyla): a review and key directions for future research. European Journal of Wildlife Research 65.

Bidder, O. R., J. S. Walker, M. W. Jones, M. D. Holton, P. Urge, D. M. Scantlebury, N. J. Marks, E. A. Magowan, I. E. Maguire, and R. P. Wilson. 2015. Step by step: reconstruction of terrestrial animal movement paths by dead-reckoning. Mov Ecol 3:23.

Camilo-Alves, C. d. S. e. P., and G. Mourao. 2005. Responses of a Specialized Insectivorous Mammal (Myrmecophaga tridactyla) to Variation in Ambient Temperature. Biotropica 38:52–56.

Díaz, S., J. Settele, E. S. Brondízio, H. T. Ngo, J. Agard, A. Arneth, P. Balvanera, K. A. Brauman, S. H. M. Butchart, K. M. A. Chan, L. A. Garibaldi, K. Ichii, J. Liu, S. M. Subramanian, G. F. Midgley, P. Miloslavich, Z. Molnár, D. Obura, A. Pfaff, S. Polasky, A. Purvis, J. Razzaque, B. Reyers, R. R. Chowdhury, Y.-J. Shin, I. Visseren-Hamakers, K. J. Willis, and C. N. Zayas. 2019. Pervasive human-driven decline of life on Earth points to the need for transformative change. Science 366:eaax3100.

Dickie, M., S. R. McNay, G. D. Sutherland, M. Cody, and T. Avgar. 2020. Corridors or risk? Movement along, and use of, linear features varies predictably among large mammal predator and prey species. J Anim Ecol 89:623–634.

Etherington, T. R. 2016. Least-Cost Modelling and Landscape Ecology: Concepts, Applications, and Opportunities. Current Landscape Ecology Reports 1:40–53.

Fagan, W. F., M. A. Lewis, M. Auger-Methe, T. Avgar, S. Benhamou, G. Breed, L. LaDage, U. E. Schlagel, W. Tang, Y. P. Papastamatiou, J. Forester, and T. Mueller. 2013. Spatial memory and animal movement. Ecology Letters 16:1316–1329.

Fletcher Jr., R. J., N. S. Burrell, B. E. Reichert, D. Vasudev, and J. D. Austin. 2016. Divergent Perspectives on Landscape Connectivity Reveal Consistent Effects from Genes to Communities. Current Landscape Ecology Reports 1:67–79.

Fletcher, R. J., and M. J. Fortin. 2018. Spatial ecology and conservation modeling: applications with R. Springer Publishing.

Fletcher, R. J., Jr., J. A. Sefair, C. Wang, C. L. Poli, T. A. H. Smith, E. M. Bruna, R. D. Holt, M. Barfield, A. J. Marx, and M. A. Acevedo. 2019. Towards a unified framework for connectivity that disentangles movement and mortality in space and time. Ecol Lett 22:1680–1689.

Forester, J. D., H. K. Im, and P. J. Rathouz. 2009. Accounting for animal movement in estimation of resource selection functions: sampling and data analysis. Ecology 90:3554–3565.

Fortin, D., H. L. Hawthorne, M. S. Boyce, D. W. Smith, T. Duchesne, and J. S. Mao. 2005. Wolves influence elk movements: behavior shapes a trophic cascade in Yellowstone National Park. Ecology 86:1320–1330.

Giroux, A., Z. Ortega, A. Bertassoni, A. L. J. Desbiez, D. Kluyber, G. F. Massocato, G. De Miranda, G. Mourao, L. Surita, N. Attias, R. d. C. Bianchi, V. P. d. O. Gasparotto, and L. G. R. Oliveira-Santos. 2021a. The role of environmental temperature on movement patterns of giant anteaters. Integrative Zoology:1–12.

Giroux, A., Z. Ortega, L. G. R. Oliveira-Santos, N. Attias, A. Bertassoni, and A. L. J. Desbiez. 2021b. Sexual, allometric and forest cover effects on giant anteaters’ movement ecology. PLOS One 16.

Gunner, R. M., M. D. Holton, D. M. Scantlebury, P. Hopkins, E. L. C. Shepard, A. J. Fell, B. Garde, F. Quintana, A. Gomez-Laich, K. Yoda, T. Yamamoto, H. English, S. Ferreira, D. Govender, P. Viljoen, A. Bruns, O. L. van Schalkwyk, N. C. Cole, V. Tatayah, L. Borger, J. Redcliffe, S. H. Bell, N. J. Marks, N. C. Bennett, M. H. Tonini, H. J. Williams, C. M. Duarte, M. C. van Rooyen, M. F. Bertelsen, C. J. Tambling, and R. P. Wilson. 2021. How often should dead-reckoned animal movement paths be corrected for drift? Anim Biotelemetry 9:43.

Haddad, N. M., L. A. Brudvig, J. Clobert, K. F. Davies, A. Gonzales, R. D. Holt, T. Lovejoy, J. O. Sexton, M. P. Austin, C. D. Collins, W. M. Cook, E. I. Damschen, R. M. Ewers, B. L. Foster, C. N. Jenkins, A. J. King, W. F. Laurance, D. J. Levey, C. R. Margules, B. A. Melbourne, A. O. Nicholls, J. L. Orrock, D.-X. Song, and J. R. Townshend. 2015. Habitat fragmentation and its lasting impact on Earth’s ecosystems. Scientific Advances 1:e1500052.

Hanks, E. M., and M. B. Hooten. 2013. Circuit Theory and Model-Based Inference for Landscape Connectivity. Journal of the American Statistical Association 108:22–33.

Harrison, S., and E. Bruna. 1999. Habitat fragmentation and large-scale conservation: what do we know for sure? Ecography 22:225–232.

Heller, N. E., and E. S. Zavaleta. 2009. Biodiversity management in the face of climate change: a review of 22 years of recommendations. Biological Conservation 142:14–32.

Hooten, M. B. 2017. Animal movement: statistical models for telemetry data. CRC Press/Taylor & Francis Group, Boca Raton.

Horne, J. S., E. O. Garton, S. M. Krone, and J. S. Lewis. 2007. Analyzing animal movements using Brownian bridges. Ecology 88:2354–2363.

Iezzi, M. E., M. S. Di Bitetti, J. M. Pardo, A. Paviolo, P. Cruz, and C. De Angelo. 2022. Forest fragments prioritization based on their connectivity contribution for multiple Atlantic Forest mammals. Biological Conservation 266.

Johnson, D. S., J. M. London, M.-A. Lea, and J. W. Durban. 2008. Continuous-Time Correlated Random Walk Model for Animal Telemetry Data. Ecology 89:1208–1215.

Kluyber, D., N. Attias, M. H. Alves, A. C. Alves, G. Massocato, and A. L. J. Desbiez. 2021. Physical capture and chemical immobilization procedures for a mammal with singular anatomy: the giant anteater (Myrmecophaga tridactyla). European Journal of Wildlife Research 67.

Kuefler, D., B. Hudgens, N. M. Haddad, W. F. Morris, and N. Thurgate. 2010. The conflicting role of matrix habitats as conduits and barriers for dispersal. Ecology 91:944–950.

Kumar, S. U., and S. A. Cushman. 2022. Connectivity modelling in conservation science: a comparative evaluation. Scientific Reports 12.

Langrock, R., R. King, J. Matthiopoulos, L. Thomas, D. Fortin, and J. M. Morales. 2012. Flexible and practical modeling of animal telemetry data: hidden Markov models and extensions. Ecology 93:2336–2342.

LaPoint, S., P. Gallery, M. Wikelski, and R. Kays. 2013. Animal behavior, cost-based corridor models, and real corridors. Landscape Ecology 28:1615–1630.

Lehnen, S. E., M. A. Sternberg, H. M. Swarts, and S. E. Sesnie. 2021. Evaluating population connectivity and targeting conservation action for an endangered cat. Ecosphere 12:e03367. 03310.01002/ecs03362.03367.

Manly, B. F. J., L. L. McDonald, D. L. Thomas, T. L. McDonald, and W. P. Erickson. 2002. Resource selection by animals: statistical design and analysis for field studies. Kluwer Academic Publishers, The Netherlands.

McDonald, T. L., B. F. J. Manly, R. M. Nielson, and L. V. Diller. 2006. Discrete-choice modeling in wildlife studies exemplified by Northern spotted owl nighttime habitat selection. The Journal of Wildlife Management 70:375–383.

McRae, B. H., B. G. Dickson, T. H. Keitt, and V. B. Shah. 2008. Using circuit theory to model connectivity in ecology, evolution, and conservation. Ecology 89:2712–2724.

Michelot, T., P. G. Blackwell, and J. Matthiopoulos. 2019. Linking resource selection and step selection models for habitat preferences in animals. Ecology 100:e02452.

Mourao, G., and I. M. Medri. 2007. Activity of a specialized insectivorous mammal (Myrmecophaga tridactyla) in the Pantanal of Brazil. Journal of Zoology 271.2:187–192.

Munden, R., L. Borger, R. P. Wilson, J. Redcliffe, R. Brown, M. Garel, and J. R. Potts. 2021. Why did the animal turn? Time-varying step selection analysis for inference between observed turning-points in high frequency data. Methods Ecol Evol 12:921–932.

Nielson, R. M., B. F. J. Manly, A. McDonald, H. Sawyer, and T. L. McDonald. 2009. Estimating habitat selection when GPS fix success is less than 100%. Ecology 90:2956–2962.

Northrup, J. M., E. V. Wal, M. Bonar, J. Fieberg, M. P. Laforge, M. Leclerc, C. M. Prokopenko, and B. D. Gerber. 2022. Conceptual and methodological advances in habitat-selection modeling: guidelines for ecology and evolution. Ecological Applications 32:e02470.

Oliveira-Santos, L. G. R., J. D. Forester, U. Piovezan, W. M. Tomas, and F. A. S. Fernandez. 2016. Incorporating animal spatial memory in step selection functions. Journal of Animal Ecology 85:516–524.

Patterson, T. A., A. Parton, R. Langrock, P. G. Blackwell, L. Thomas, and R. King. 2017. Statistical modelling of individual animal movement: an overview of key methods and a discussion of practical challenges. AStA Advances in Statistical Analysis 101:399–438.

Plummer, M. 2003. JAGS: A program for analysis of Bayesian graphical models using GIbbs sampling.

Pohle, J., R. Langrock, F. M. van Beest, and N. M. Schmidt. 2017. Selecting the number of states in hidden Markov models: pragmatic solutions illustrated using animal movement. Journal of Agricultural, Biological, and Environmental Statistics 22:270–293.

Potts, J. R., G. Bastille-Rousseau, D. L. Murray, J. A. Schaefer, and M. A. Lewis. 2014. Predicting local and non-local effects of resources on animal space use using a mechanistic step selection model. Methods Ecol Evol 5:253–262.

Prevedello, J. A., G. Forero-Medina, and M. V. Vieira. 2010. Movement behaviour within and beyond perceptual ranges in three small mammals: effects of matrix type and body mass: Movement behaviour and perceptual range. Journal of Animal Ecology 79:1315–1323.

Prokopenko, C. M., M. S. Boyce, and T. Avgar. 2017. Characterizing wildlife behavioural responses to roads using integrated step selection analysis. Journal of Applied Ecology 54:470–479.

Searle, K. R., N. T. Hobbs, and L. A. Shipley. 2005. Should I stay or should I go? Patch departure decisions by herbivores at multiple scales. Oikos 111:417–424.

Semenchuk, P., C. Plutzar, T. Kastner, S. Matej, G. Bidoglio, K.-H. Erb, F. Essl, H. Haberl, J. Wessely, F. Krausmann, and S. Dullinger. 2022. Relative effects of land conversion and land-use intensity on terrestrial vertebrate diversity. Nature Communications 615.

Van Moorter, B., C. M. Rolandsen, M. Basille, and J.-M. Gaillard. 2016. Movement is the glue connecting home ranges and habitat selection. Journal of Animal Ecology 85:21–31.

Vardakis, M., P. Goos, F. Adriaensen, E. Matthysen, and J. Matthiopoulos. 2015. Discrete choice modelling of natal dispersal: ‘Choosing’ where to breed from a finite set of available areas. Methods in Ecology and Evolution 6:997–1006.

Zeller, K. A., K. McGarical, and A. R. Whiteley. 2012. Estimating landscape resistance to movement: a review. Landscape Ecology 27:777–797.

Zeller, K. A., K. McGarigal, S. A. Cushman, P. Beier, T. W. Vickers, and W. M. Boyce. 2016. Using step and path selection functions for estimating resistance to movement: pumas as a case study. Landscape Ecology 31:1319–1335.

Zollner, P. A., and S. L. Lima. 1999. Search Strategies for Landscape-Level Interpatch Movements. Ecology 80:1019–1030.

