## Appendix 1 for "Bridging the gap between movement data and connectivity analysis using the time-explicit Step Selection Function (tSSF)"

### Appendix 1. Simulations for the time model

We rely on simulations to determine how well the time model performs. More specifically, recall that the time model relies on an approximation of the gamma process in which averaged covariates are used within each step. We expect that this approximation will work well if the area traversed within each step is relatively homogeneous. This is more likely to be true if the landscape is relatively homogeneous, if the time interval between GPS fixes is short, and/or the animal does not move very fast (i.e., the distance traversed is small).

We simulated 1-D landscapes with 1,000,000 cells and 3 possible land-use/land-cover (LULC) classes. We populate this 1-D landscape by iteratively sampling the LULC class and assuming that $N$ neighboring pixels had this class. For example, if we sampled LULC classes 1, 3, and 1 and $N$ was equal to 3, that would mean that our landscape would be comprised of the following sequence of pixels classified according their LULC class [1,1,1,3,3,3,1,1,1]. To make the landscape more and more homogenous within each step, we assumed that N was equal to 1, 10, 100 and 1,000. Figs. S1A and S1B illustrate that, as N increases and the corresponding landscape becomes more homogeneous (top to bottom panels), the average proportion of each LULC within each step tends to become almost exclusively comprised of 0's and 1's.

We set the intercept to -1 and the slopes associated with the second and third LULC classes to 0.1 and -0.3, respectively. Notice that we do not include the first LULC class because this class serves as the baseline class in our model. The over-dispersion parameter b was set to 2. Based on these parameter values, we randomly sampled the amount of time (in minutes) taken to traverse each pixel by drawing from a gamma distribution. Finally, we assume that the time between fixes was approximately 10 minutes. As a result, we simulated the animal moving through this 1-D landscape and each step was defined as the set of pixels that were traversed within approx. 10 minutes. Within each step, we calculated the number of traversed pixels (i.e., the traversed distance) and the proportion of each LULC classes among the traversed pixels.

Our results reveal that the proposed time model, despite approximating $\sum_{i=1}^{n_{j}} D_{ij}\exp\left( \boldsymbol{x}_{\boldsymbol{i}}^{\boldsymbol{T}}\boldsymbol{\beta} \right)$ by $D_{j}\exp\left( {\bar{\boldsymbol{x}}}_{\boldsymbol{j}}\boldsymbol{\beta} \right)$, can still estimate well all the parameters in the model (Fig. S1c). Nevertheless, although it is hard to see in Fig. S1c, there tends to be greater parameter uncertainty (as captured by the width of the 95% credible intervals) as landscape heterogeneity within each step increases.

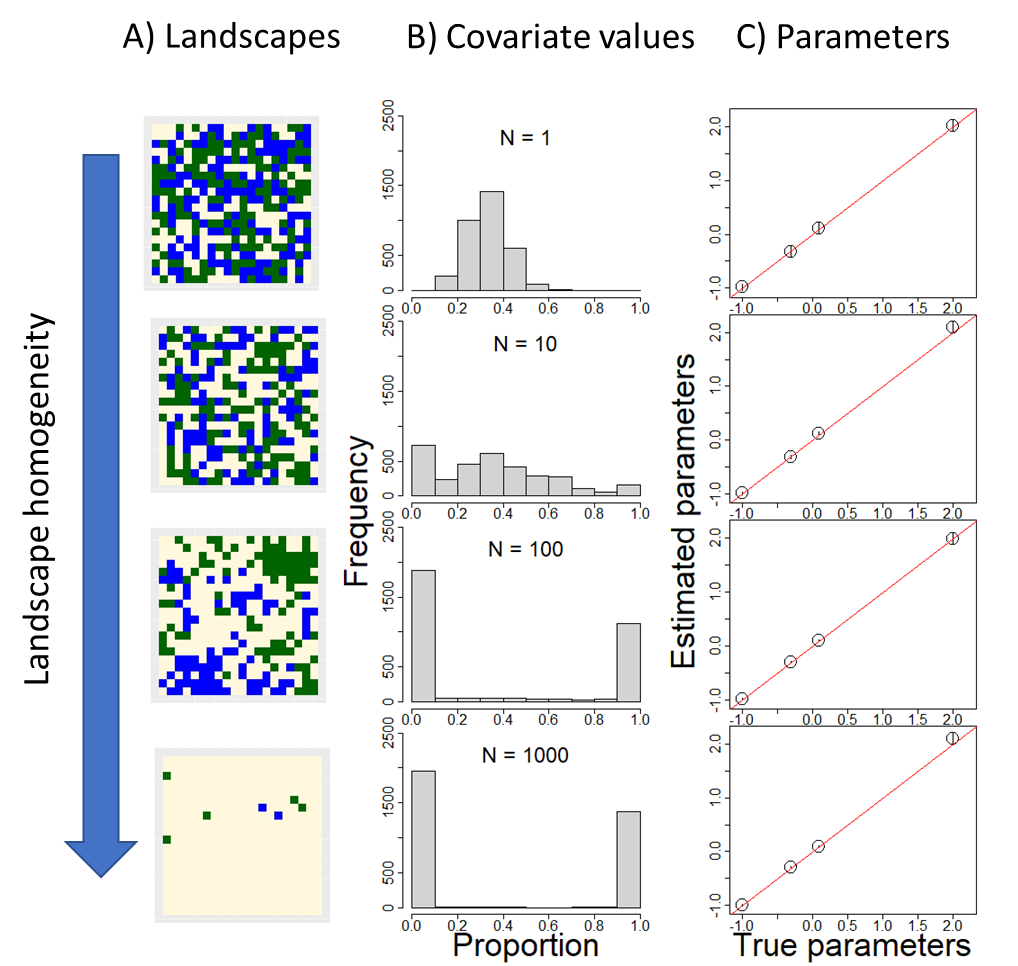

Fig. S1. Results for the simulated data, where panels are ordered top to bottom from more heterogeneous to more homogeneous landscapes. (a) These panels illustrate the different types of landscapes (each color represents a different LULC class). (b) Distribution of the proportion of a LULC class within each step for each hypothetical landscape. N represents the number of neighboring cells that contain the same LULC class and is a proxy of landscape homogeneity (i.e., the higher the value of N, the more homogeneous the landscape is). (c) These panels compare the estimated to the true parameter values, where the 1:1 line is shown in red. Vertical black lines in panel C are 95% credible intervals but are hard to see because they are very short.
