## Appendix 2 for "Bridging the gap between movement data and connectivity analysis using the time-explicit Step Selection Function (tSSF)"

### Appendix 2. Simulations for the time-integrated Step Selection function (tSSF)

We rely on simulations to evaluate how inference on step selection strength is influenced: by a) the number of available steps used in the model; and b) how these available steps are selected. In relation to the number of available steps, we compared model results assuming one versus four available steps for each observed step. In relation to how available steps are selected, we fit the model to data where potential step lengths are chosen at random from the empirical distribution of step lengths, as traditionally done in step selection models, and we compare these modeling results to those obtained by pairing each observed step with potential steps of similar length. In total, we compare the results from four different scenarios that arise from combining two different approaches to selecting available steps with two different numbers of available steps (i.e., 1 vs. 4 available steps).

For each simulation, we assume that 1000 steps were observed. For each step, we generate a landscape surrounding the starting point consisting of 900 pixels with 3 land-use/land-cover (LULC) classes. We then calculate the proportion of each LULC for all possible steps (i.e., all steps starting at the center cell to each of the 900 pixels that comprise the landscape; grey lines in Fig. S1). We adopted the same parameters for the time model as the ones used in Appendix 1. In other words, the intercept was set to -1, the first LULC class was used as the baseline class, the slopes associated with the second and third LULC classes were set to 0.1 and -0.3, respectively, and the over-dispersion parameter b was set to 2. However, because each landscape is relatively small, we assumed that the time interval between GPS fixes is equal to 2 min (in contrast to the 20-30 min intervals of our case study).

In relation to the tSSF, we set the slope for the second and third LULC classes to 0.5 and -0.1, respectively. Using these parameter values and the LULC proportion for each possible step, our simulation draws the actual step taken by the animal (red line in Fig. S1) from a categorical distribution having one probability associated with each of the 900 potential steps. These probabilities are calculated based on the tSSF equation described in the main document.

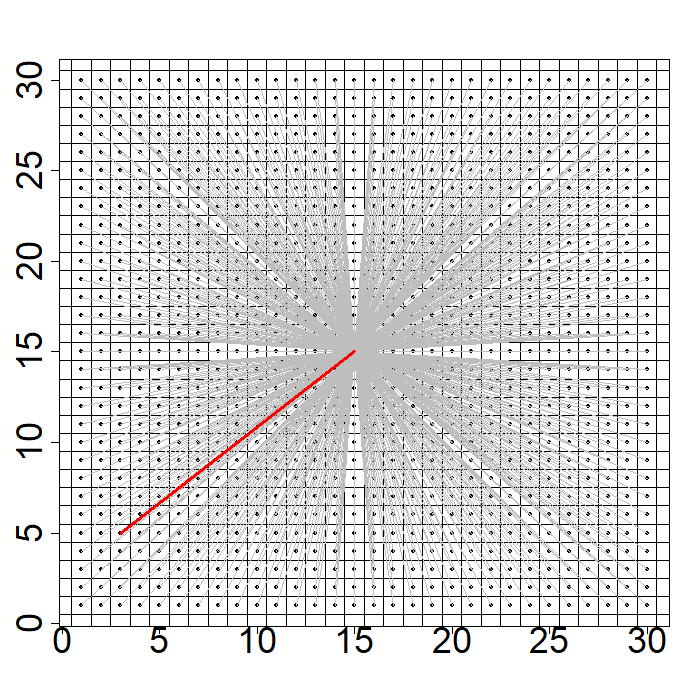

Fig. S1 Illustration of all of the 900 available steps that start from the center cell (grey lines). The actual step taken by the animal is depicted with the red line.

As expected, parameter estimates were unbiased for all scenarios, regardless of the number of available steps and how these steps were selected. However, adding more available steps tended to result in increased precision in parameter estimates (i.e., narrower 95% credible intervals; Fig. S2). Furthermore, as expected, we did not see major differences between selecting from the empirical distribution of steps lengths versus using step lengths of similar size to the paired observed step.

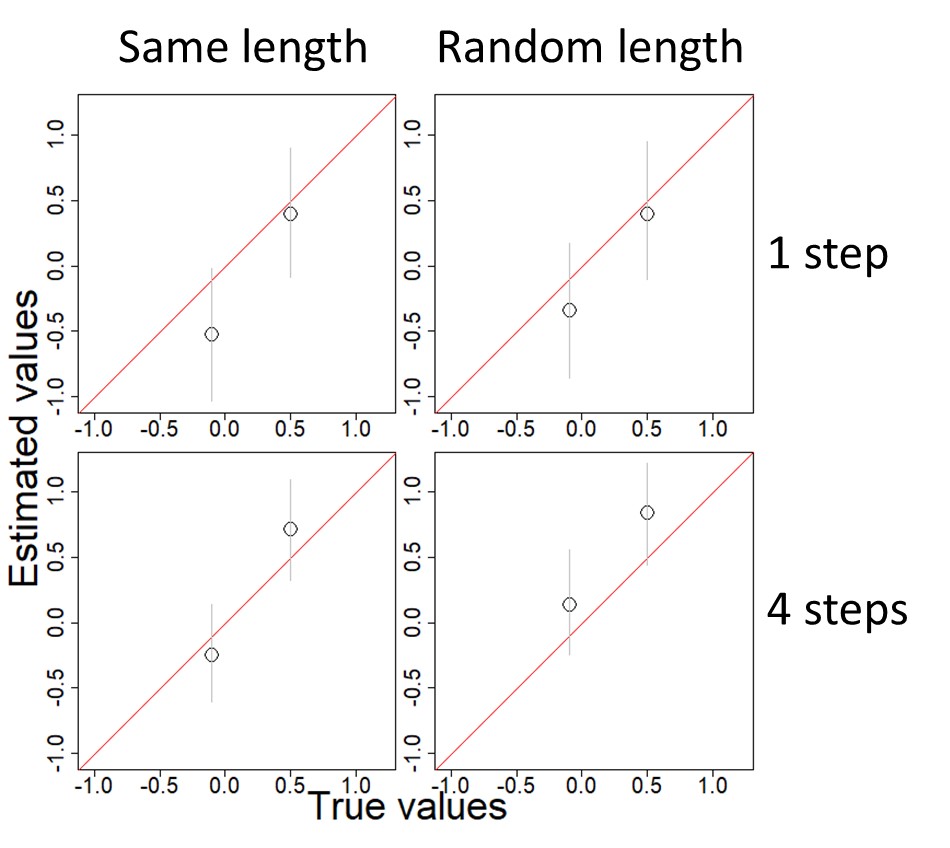

Fig. S2. Comparison of parameter estimates and their true values for tSSF using simulated data for four different scenarios. The posterior median (black circles) and 95% credible intervals (grey vertical lines) are shown for models fit to data from each of these different scenarios. For two datasets, the length of available steps was determined by sampling from the empirical distribution of step lengths ("random length", right panels). For two datasets, available steps were chosen by sampling steps with the same length as the paired observed step ("same length", left panels). Results are also shown for models using data with 1 ("1 step", top panels) and 4 ("4 steps", bottom panels) available steps paired to each observed step. A 1:1 red line is shown for reference.
