## Appendix 4 for "Bridging the gap between movement data and connectivity analysis using the time-explicit Step Selection Function (tSSF)"

### Appendix 4. Results associated with $\gamma_{1}$ in the time model

Here we report the results for the parameter $\gamma_{1}$ from the time model applied to the giant anteater case study. Recall that this parameter quantifies if the time taken to traverse a particular area has greater uncertainty when one or more GPS fixes are missed. All of the 99% credible intervals for $\gamma_{1}$ did not overlap zero and the posterior medians were negative (Table S1), revealing that observations with missed GPS fixes tend to have greater uncertainty. This is expected given that there is less information regarding the actual path taken by the animal within that timeframe.

Table 1. Posterior summaries for $\gamma_{1}$.

| ID | Median | 99% credible interval | |
| --- | --- | --- | --- |
|  |  | 0.5% | 99.5% |
| Berenice | -0.84 | -1.01 | -0.66 |
| Brigite | -0.46 | -0.58 | -0.35 |
| Fergus | -0.76 | -1.02 | -0.54 |
