## Appendix 5 for "Bridging the gap between movement data and connectivity analysis using the time-explicit Step Selection Function (tSSF)"

Appendix 6. Nonsensical parameter estimates associated with the movement kernel within iSSF.

Goal:

Using simulations, we show how fitting iSSF using a gamma distribution for its movement kernel can generate negative shape and scale parameters even if the data generating mechanism is based on positive parameter values.

Description of the iSSF model

Assume a gamma distribution for the movement kernel, given by:

$$p\left( y_{ij} \right)=\frac{\left( \frac{1}{q} \right)^{k}}{\Gamma\left( k \right)}y_{ij}^{k-1}exp(-\frac{1}{q}y_{ij})$$

where $y_{ij}$ is the step-length (i.e., the distance between locations i and j) and k and q are the shape and scale parameters respectively. For simplicity, we ignore resource selection and the distribution for turning angles, enabling us to write the iSSF model as:

$$p\left( P_{t+\Delta t}=j|\Delta t,P_{t}=i \right)=\frac{\exp\left( -\frac{1}{q}y_{ij}+\left( k-1 \right)\log\left( y_{ij} \right) \right)}{\sum_{w} \exp\left( -\frac{1}{q}y_{iw}+\left( k-1 \right)\log\left( y_{iw} \right) \right)}$$

where $P_{t}$ and $P_{t+\Delta t}$ are the locations at time t and time $t+\Delta t$, respectively.

Let $-\frac{1}{q}=\beta_{1}$ and $k-1=\beta_{2}$. Furthermore, assume that $\beta_{1}$ and $\beta_{2}$ are linear functions of covariate $x_{ij}\in\left[ 0,1 \right]$ (e.g., the proportion of a LULC class along the path between locations i and j). In other words,

$$\beta_{1ij}=\theta_{1}+\theta_{2}x_{ij}$$

$$\beta_{2ij}=\theta_{3}+\theta_{4}x_{ij}$$

As a result, we can write our original expression as

$$p\left( P_{t+\Delta t}=j|\Delta t,P_{t}=i \right)=\frac{\exp\left( \left[ \theta_{1}+\theta_{2}x_{ij} \right]y_{ij}+\left[ \theta_{3}+\theta_{4}x_{ij} \right]\log\left( y_{ij} \right) \right)}{\sum_{w} \exp\left( \left[ \theta_{1}+\theta_{2}x_{iw} \right]y_{iw}+\left[ \theta_{3}+\theta_{4}x_{iw} \right]\log\left( y_{iw} \right) \right)}$$

In relation to our constraints, we know that $q>0$ and therefore $\beta_{1ij}=\theta_{1}+\theta_{2}x_{ij}<0$. Furthermore, because $x_{ij}\in\left[ 0,1 \right]$, this implies that $\theta_{1}<0$ and $\theta_{1}+\theta_{2}<0$. Similarly, because $k>0$, this implies that $\beta_{2ij}=\theta_{3}+\theta_{4}x_{ij}>-1$. Furthermore, because $x_{ij}\in\left[ 0,1 \right]$, this implies that $\theta_{3}>-1$ and $\theta_{3}+\theta_{4}>-1$. In short, the natural constraints in the parameters of the gamma distribution imply the following constraints on these model parameters:

$$\theta_{1}<0$$

$$\theta_{1}+\theta_{2}<0$$

$$\theta_{3}>-1$$

$$\theta_{3}+\theta_{4}>-1$$

Simulation approach for the iSSF model

We simulate data for iSSF by sampling from:

$$p\left( P_{t+\Delta t}=j|\Delta t,P_{t}=i \right)=\frac{\exp\left( \left[ \theta_{1}+\theta_{2}x_{ij} \right]y_{ij}+\left[ \theta_{3}+\theta_{4}x_{ij} \right]\log\left( y_{ij} \right) \right)}{\sum_{w} \exp\left( \left[ \theta_{1}+\theta_{2}x_{iw} \right]y_{iw}+\left[ \theta_{3}+\theta_{4}x_{iw} \right]\log\left( y_{iw} \right) \right)}$$

We created artificial landscapes containing 900 pixels, with the individual starting in the middle of these landscapes. We randomly drew the location chosen by the individual from a categorical distribution with probabilities governed by the iSSF probabilities. Finally, we kept the chosen pixel and randomly chose 4 additional pixels that were available but that were not chosen. On total, we simulated 1,000 of these landscapes and thus each iSSF dataset contained (1 selected + 4 available pixels) x 1,000 landscapes = 5,000 observations.

To satisfy the constraints on parameter values when simulating the data, we drew these parameters from the following distributions:

$$\theta_{1}\sim Unif\left( -1,0 \right)$$

$$\theta_{2}|\theta_{1}\sim Unif\left( -1,-\theta_{1} \right)$$

$$\theta_{3}\sim Unif\left( -1,1 \right)$$

$$\theta_{4}|\theta_{3}\sim Unif\left( -1-\theta_{3},1 \right)$$

On total, 100 simulated datasets were created. The iSSF model was fitted within a Bayesian framework using JAGS, using code that is similar to the one used for the tSSF model.

Results for iSSF

Our results reveal that the model parameters were well estimated for this iSSF model, with a comparison between the estimated and the true parameters generally falling along the 1:1 line (Fig. 1). However, despite the fact that the simulations were created with parameter values that satisfied the constraints imposed by the gamma distribution assumption for step-lengths, some parameter estimates violated these constraints (red circles in Fig. 1). Furthermore, the resulting posterior distributions often placed considerable weight on nonsensical parameter values, as evident by some of the 95% credible intervals violating these parameter constraints (red vertical lines in Fig. 1). The reason for this is because the iSSF, as currently formulated, places no restriction on parameter values.

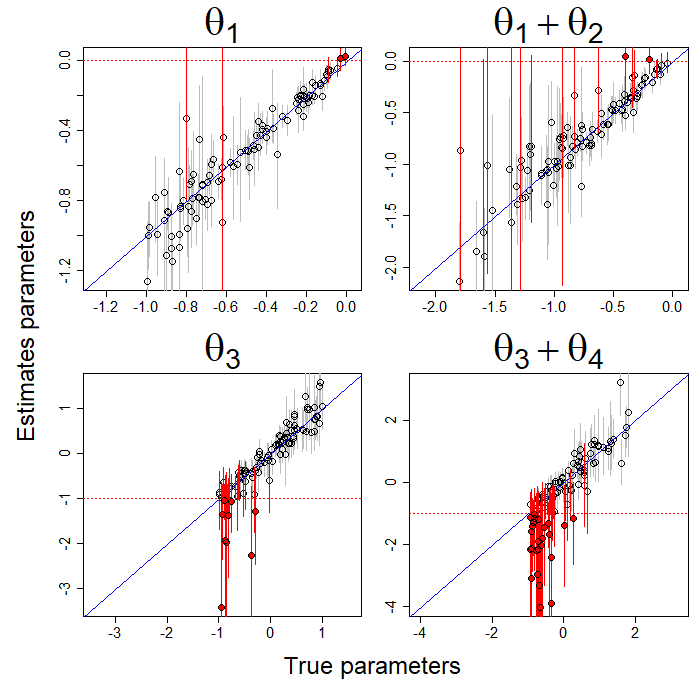

Fig. 1. Comparison of the true and estimated parameter values. A 1:1 line (blue diagonal line) was added for reference. Dotted horizontal red lines represent the constraint value for each parameter/set of parameters. Each circle represents the result for one of the simulated data sets. Red and black circles are results that violate and that do not violate the constraint, respectively. The 95% credible intervals are depicted in red and grey when they overlap and do not overlap the constraint value, respectively.
