## Appendix 6 for "Bridging the gap between movement data and connectivity analysis using the time-explicit Step Selection Function (tSSF)"

Appendix 7. Parameter identifiability associated with the iSSF model.

Goal:

Using simulations, we show that fitting iSSF using a gamma distribution for its movement kernel can be challenging given the high-correlation induced by including step-lengths, log(step-lengths), and their interaction with other covariates.

Description of the iSSF model

To simulate data for iSSF, we assume a gamma distribution for step-lengths. Furthermore, we assume that both parameters of the gamma distribution (shape and scale) are functions of two covariates $\bar{x}_{1ij}$ and $\bar{x}_{2ij}$. Similarly, we assume that selection is also a function of these two covariates. Notice that we do not include a distribution for turning angles. This model is given by:

$$p\left( P_{t+\Delta t}=j|\Delta t,P_{t}=i \right)=\frac{\exp\left( \beta_{0}\bar{x}_{1ij}+\beta_{1}\bar{x}_{2ij}+\beta_{2}y_{ij}+\beta_{3}\log\left( y_{ij} \right)+\beta_{4}\bar{x}_{1ij}\log\left( y_{ij} \right)+\beta_{5}\bar{x}_{2ij}\log\left( y_{ij} \right)+\beta_{6}\bar{x}_{1ij}y_{ij}+\beta_{7}\bar{x}_{2ij}y_{ij} \right)}{\sum_{k} \exp\left( \beta_{0}\bar{x}_{1ik}+\beta_{1}\bar{x}_{2ik}+\beta_{2}y_{ik}+\beta_{3}\log\left( y_{ik} \right)+\beta_{4}\bar{x}_{1ik}\log\left( y_{ik} \right)+\beta_{5}\bar{x}_{2ik}\log\left( y_{ik} \right)+\beta_{6}\bar{x}_{1ik}y_{ik}+\beta_{7}\bar{x}_{2ik}y_{ik} \right)}$$

where $y_{ij}$ is the distance between pixel i and pixel j (i.e., step-length).

We created artificial landscapes containing 900 pixels, with the individual starting in the middle of these landscapes. We randomly drew the location chosen by the individual from a categorical distribution with probabilities governed by the iSSF probabilities. Finally, we kept the chosen pixel and randomly chose 4 additional pixels that were available but that were not chosen. On total, we simulated 1,000 of these landscapes and thus each iSSF dataset contained (1 selected + 4 available pixels) x 1,000 landscapes = 5,000 observations.

On total, 100 simulated datasets were created. Parameters $\beta_{0},\ldots,\beta_{7}$ were randomly drawn from a uniform distribution between -1 and 1. The iSSF model was fitted both in a Bayesian framework (using JAGS, with code that is similar to the one used for the tSSF model) and in a maximum likelihood framework (using the function “clogit” within the R package “survival”).

Results for iSSF

Our results using a Bayesian framework reveal that some of the iSSF model parameters were well estimated, with a comparison between the estimated and the true parameters generally falling along the 1:1 line, whereas other parameters were less well estimated (Fig. 1). Importantly, we find that 22% of the models fitted to the 100 simulated datasets had at least one parameter that did not converge (i.e., convergence statistic $\hat{R}$>1.1). Table 1 describes the proportion of times that each parameter had $\hat{R}$>1.1. This is likely a result of the high correlation between $y_{ij}$ and $\log\left( y_{ij} \right)$, which induces a high correlation (r>0.9) between the following pairs of parameters: $\beta_{2}$ and $\beta_{3}$, $\beta_{4}$ and $\beta_{6}$, and $\beta_{5}$ and $\beta_{7}$.

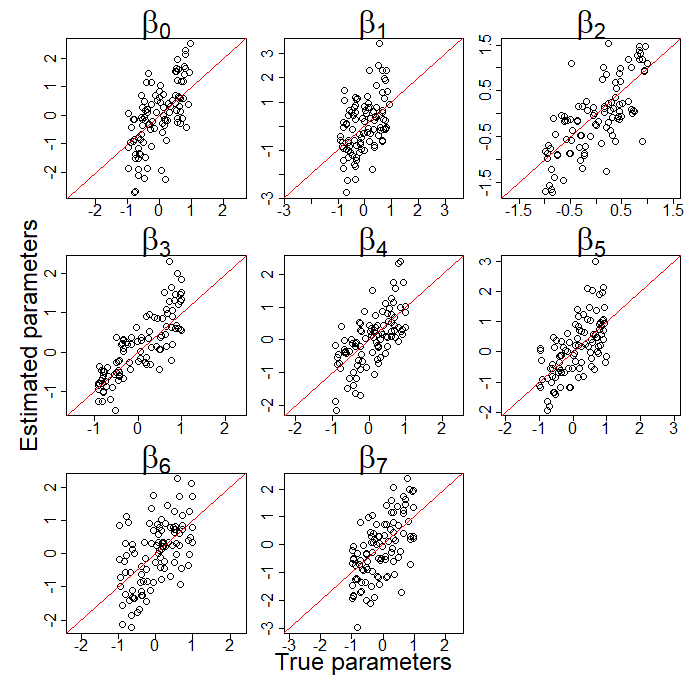

Fig. 1. Comparison of the estimated and true parameter values for each iSSF parameter. Parameters were estimated in a Bayesian framework. A 1:1 line was added for reference (diagonal red line). Each circle represents the result based on one of the 100 simulated datasets.

| Parameter | Proportion with lack of convergence |
| --- | --- |
| $\beta_{0}$ | 0.13 |
| $\beta_{1}$ | 0.13 |
| $\beta_{2}$ | 0.18 |
| $\beta_{3}$ | 0.14 |
| $\beta_{4}$ | 0.09 |
| $\beta_{5}$ | 0.09 |
| $\beta_{6}$ | 0.13 |
| $\beta_{7}$ | 0.13 |

Table 1. Proportion of times (out of 100 Bayesian models) that each parameter did not converge (i.e., convergence statistic $\hat{R}$>1.1).

Finally, we note that the correlation between $y_{ij}$ and $\log\left( y_{ij} \right)$ is problematic irrespective of the framework used to fit the model (i.e., Bayesian or MLE). For example, when these data are fitted in a maximum likelihood framework, we find a similar result to those from the Bayesian model when comparing the estimated and the true parameter values (Fig. 2). More critically however, we observe 95% confidence intervals with widths in the order of the tens to hundreds, which are very large given that the true parameter values varied from -1 and 1 (Table 2).

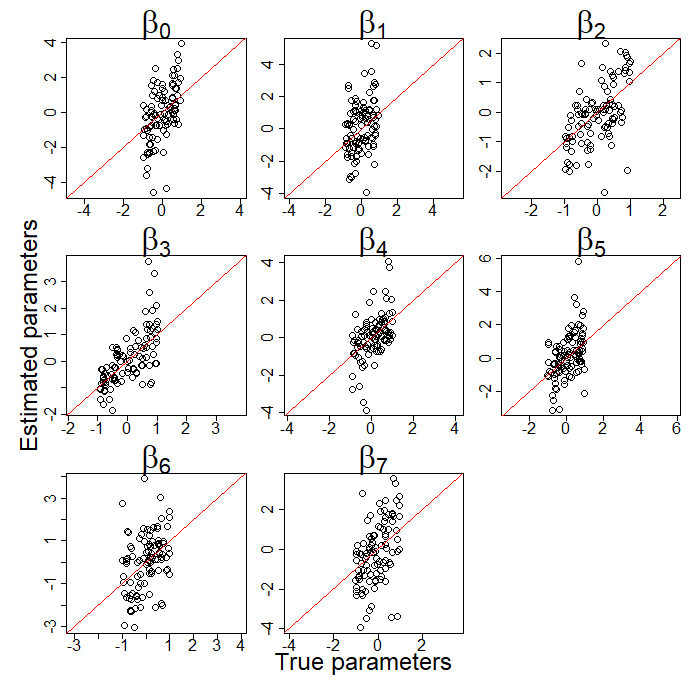

Fig. 2. Comparison of the estimated and true parameter values for each iSSF parameter. Parameters were estimated in a MLE framework using the R package “survival”. A 1:1 line was added for reference (diagonal red line). Each circle represents the result based on one of the 100 simulated datasets.

| Parameter | Mean width of 95% confidence intervals |
| --- | --- |
| $\beta_{0}$ | 48 |
| $\beta_{1}$ | 203 |
| $\beta_{2}$ | 6 |
| $\beta_{3}$ | 16 |
| $\beta_{4}$ | 83 |
| $\beta_{5}$ | 309 |
| $\beta_{6}$ | 41 |
| $\beta_{7}$ | 43 |

Table 2. Mean width of 95% confidence intervals for each parameter based on 100 simulated datasets. Parameters were estimated in a MLE framework using the R package “survival”.
