## Appendix 3 for "Bridging the gap between movement data and connectivity analysis using the time-explicit Step Selection Function (tSSF)"

### Data preparation for the time and tSSF models

Denis Valle

March 2023

#### Importing the data

The data come from a giant anteater monitored in the Brazilian Pantanal region. Each row in our data set consists of the information regarding one GPS fix. The columns in this dataset are:

- `timestamp`: day and time of GPS fix; and
- `utm.easting` and `utm.northing`: spatial coordinates of GPS fix.

We start by importing these data.

```
rm(list=ls())
library('lubridate')
library(sf)
library(raster)
library(lubridate)
library('terra')
library('rgdal')

#get data
setwd('U:\\timemod\\tutorials\\prep data tutorial')
dat1=read.csv('data0.csv')
head(dat1)
```

```
##             timestamp utm.easting utm.northing
## 1 2015-06-20 16:23:04      621193      7865263
## 2 2015-06-20 17:23:17      621113      7864660
## 3 2015-06-20 18:23:12      621194      7865267
## 4 2015-06-20 19:23:19      621093      7864997
## 5 2015-06-20 20:40:38      621188      7865279
## 6 2015-06-20 21:01:07      621199      7865261
```

#### Calculating time, distance, and Miss columns and cleaning the data

Then, we calculate step length and time interval between consecutive GPS fixes.

```
#make sure these data are ordered chronologically
dat1$timestamp=as_datetime(dat1$timestamp, tz = "UTC")
dat2=dat1[order(dat1$timestamp),]

#define starting and end points
dat2$x.sta=dat2$utm.easting   #x coordinate for starting point
dat2$y.sta=dat2$utm.northing  #y coordinate for starting point
```

```
dat3=dat2
dat3$x.end=c(dat3$utm.easting[-1],NA)  #x coordinate for end point
dat3$y.end=c(dat3$utm.northing[-1],NA) #y coordinate for end point

#get time interval
dat3$time.sta=dat3$timestamp
dat3$time.end=c(dat3$timestamp[-1],NA)
dat3$time1=as.numeric(difftime(dat3$time.end,dat3$time.sta,unit='mins'))

#get distance
x2=(dat3$x.end-dat3$x.sta)^2
y2=(dat3$y.end-dat3$y.sta)^2
dat3$dist1=sqrt(x2+y2)

#look at the distribution of interval time between GPS fixes
hist(dat3$time1[dat3$time1<70])
abline(v=35)
```

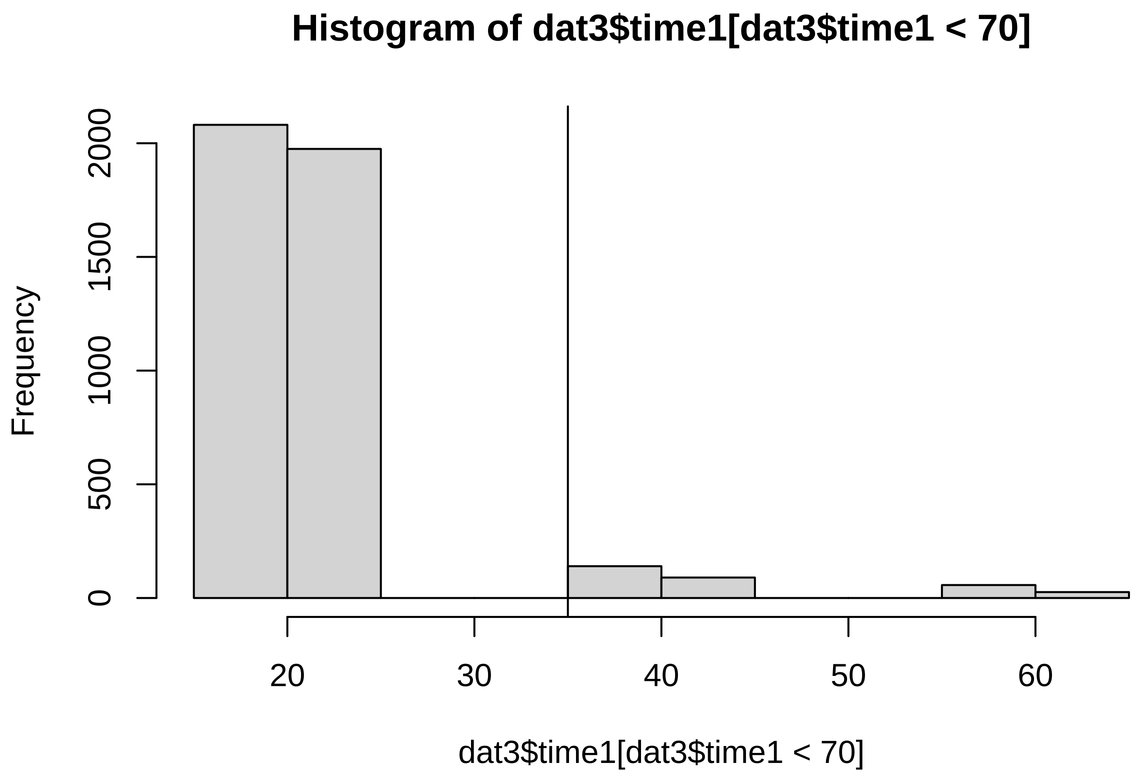


```
#create new column indicating if one or more GPS fixes were missed
dat3$Miss=ifelse(dat3$time1<35,0,1)
```

Now we clean these data by removing missing observations. Furthermore, because the time model requires that distance is different from zero, we replace all zero distances by 1 meter (or another small number). Finally, we remove paths that imply unrealistic speeds and paths for which the time interval is too long (i.e., > 60 min.).

```
#remove observations for which dist1 or time1 are missing
# apply(is.na(dat3),2,sum)
cond=is.na(dat3$dist1) | is.na(dat3$time1)
dat4=dat3[!cond,]

#if dist1==0, set to 1
cond=dat4$dist1==0; dat4$dist1[cond]=1
# mean(cond);
# sum(cond); only 4 observations

#remove extreme speeds and time intervals > 60
speed=dat4$dist1/dat4$time1

#examine distribution of speed
hist(speed)
```

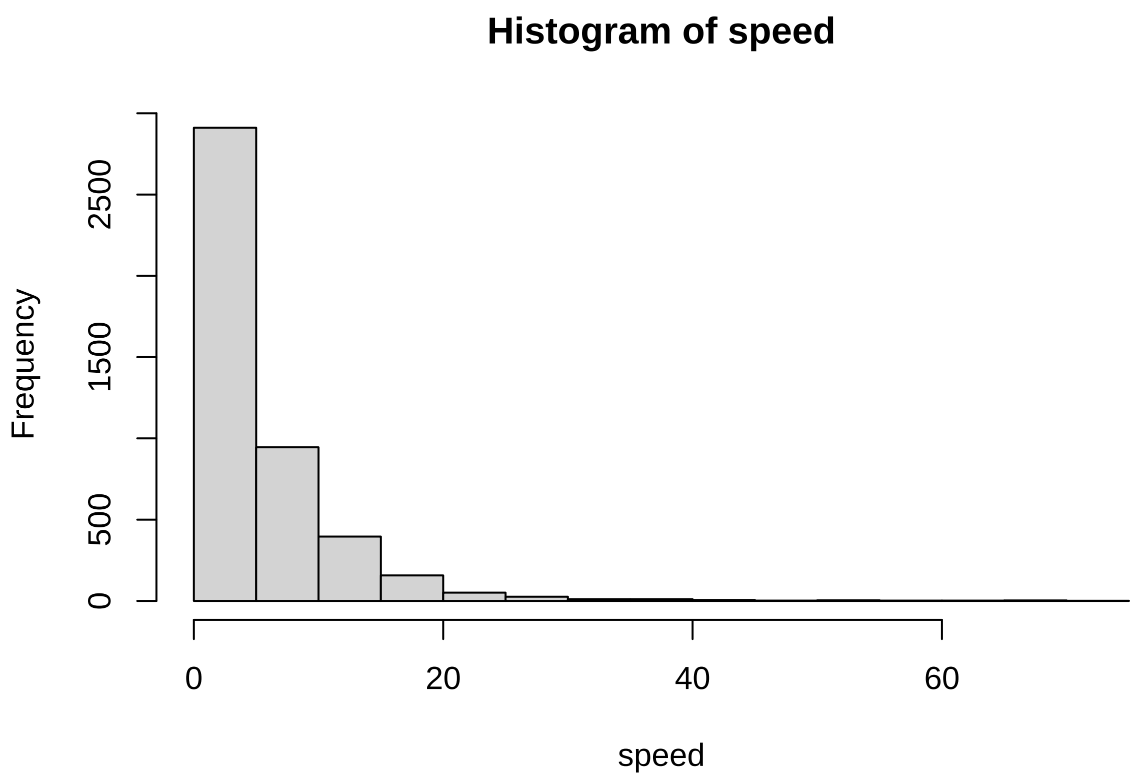


```
quantile(speed,0.99) # 28.99 m/s
```

```
##      99%
## 28.9935
```

```
#just keep observations for which speed is less than 30m and time interval is less than 60 min
cond=speed<30 & dat4$time1 <= 60; #hist(speed[cond])
dat5=dat4[cond,]
```

#### Extracting LULC information

To extract the land-use/land-cover (LULC) information along each step, we have to define the projection that our data are on and we have to import the raster file containing the LULC information.

```
#get coordinate system
EPSG=make_EPSG()
ind=grep('WGS 84 / UTM zone 21S',EPSG$note);
#EPSG$note[ind]
prj.info=EPSG[ind,'prj4']

#get LULC
setwd('U:\\timemod\\tutorials\\prep data tutorial')
ras1 <- raster('lulc subset.tif')
plot(ras1)
```

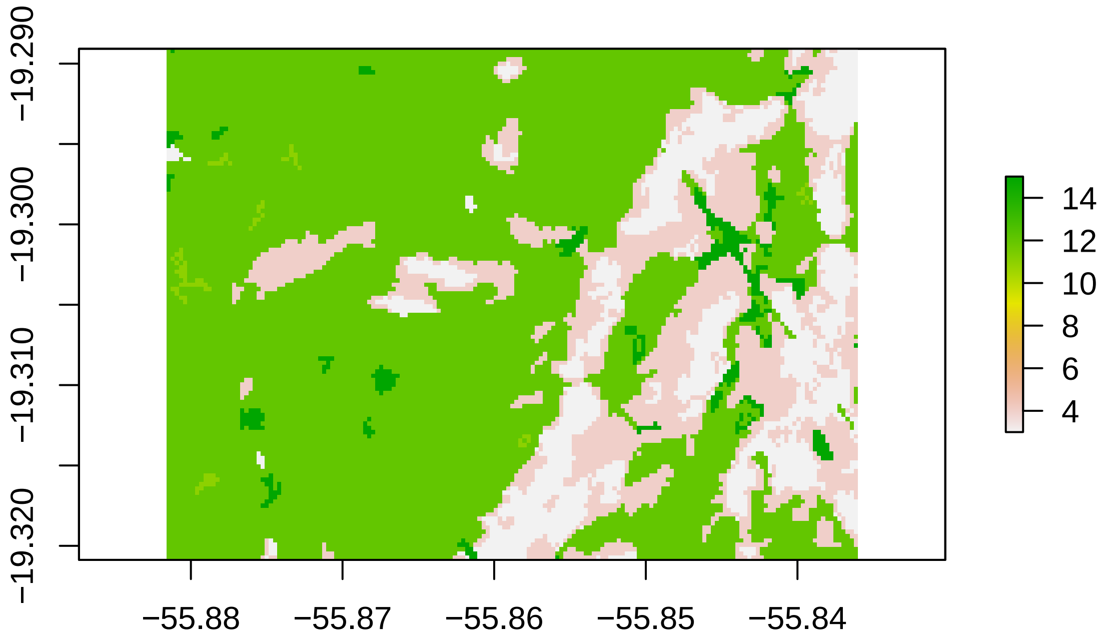


At this point, we are ready to begin extracting the LULC information. For each path, we:

a) create a line segment;
b) define its coordinate system;
c) create a 30 m buffer around the line segment;
d) extract information from the raster using our buffer;
e) tabulate the number of each type of LULC pixel and convert to proportion; and
f) store the results.

Notice that, because there are at most 50 LULC categories, we create an empty matrix containing 50 columns and use this matrix to store the results for each path.

```
#extract LULC along each path
nobs=nrow(dat5)
lulc=matrix(NA,nobs,50)
for (i in 1:nobs){
  print(i)

  #create line segment
  coord=cbind(c(dat5$x.sta[i],dat5$x.end[i]),
              c(dat5$y.sta[i],dat5$y.end[i]))
  seg1= st_linestring(coord)

  #define coordinate system
  seg2=st_sfc(seg1,crs = prj.info)
  seg3=st_sf(seg2)

  #create buffer
  seg4=st_buffer(seg3,dist=30)
  seg5=st_transform(seg4,crs=crs(ras1))

  #extract from raster
  tmp1<- raster::extract(ras1, seg5, along = TRUE, cellnumbers = F,
                         method='simple')

  #tabulate the number of each type of LULC pixel and convert to proportion
  tmp2=table(tmp1[[1]])
  tmp3=tmp2/sum(tmp2)
  tmp4=rep(0,50)
  tmp4[as.numeric(names(tmp3))]=tmp3
  lulc[i,]=tmp4
}
```

Finally, we just keep the columns of LULC categories that truly exist in our landscape and rename the columns according to what the different numbers in this raster represent.

```
#just keep values that really show up
colnames(lulc)=paste0('lulc',1:ncol(lulc))
sum1=apply(lulc,2,sum)
ind=which(sum1==0)
lulc1=lulc[,-ind]
colnames(lulc1)=paste0(c('forest','savanna','wetland','grass','pasture'),0)

#codes from mapbiomas
# 3: forest formation
# 4: savanna formation
# 11: wetland
# 12: grassland formation
# 15: pasture
# 33: water

#combine with original data
dat6=cbind(dat5[,c('time1','dist1','x.sta','y.sta','Miss')],lulc1)
head(round(dat6,2))
```

```
##   time1  dist1  x.sta   y.sta Miss forest0 savanna0 wetland0 grass0 pasture0
## 2 59.92 612.38 621113 7864660    1    0.13     0.76        0   0.09     0.02
```

```
## 5 20.48  21.10 621188 7865279    0    0.00     0.33        0   0.33     0.33
## 6 20.43  56.44 621199 7865261    0    0.00     0.29        0   0.57     0.14
## 7 39.80  15.03 621171 7865212    1    0.00     0.20        0   0.80     0.00
## 8 20.23 103.24 621172 7865227    0    0.09     0.18        0   0.64     0.09
## 9 19.68 117.69 621165 7865330    0    0.08     0.25        0   0.42     0.25
```

```
#remove columns 'x.sta' and 'y.sta' and export time model data
ind=which(colnames(dat6)%in%c('x.sta','y.sta'))
setwd('U:\\timemod\\tutorials\\prep data tutorial')
write.csv(dat6[,-ind],'timemod data1.csv',row.names=F)
```

This dataset now contains all the information that we need to fit the time model.

### Creating potential steps for tSSF model

The tSSF model, on the other hand, requires the creation of additional steps. In our work, we have relied on the creation of 4 potential steps, one in each cardinal direction and all having the same length as the realized step. Here is how we obtain the coordinates of each one of these potential steps.

```
#get step to the west
dat6$x.end1=dat6$x.sta-dat6$dist1
dat6$y.end1=dat6$y.sta

#get step to the east
dat6$x.end2=dat6$x.sta+dat6$dist1
dat6$y.end2=dat6$y.sta

#get step to the north
dat6$x.end3=dat6$x.sta
dat6$y.end3=dat6$y.sta+dat6$dist1

#get step to the south
dat6$x.end4=dat6$x.sta
dat6$y.end4=dat6$y.sta-dat6$dist1
head(round(dat6,2))
```

```
##   time1  dist1  x.sta   y.sta Miss forest0 savanna0 wetland0 grass0 pasture0
## 2 59.92 612.38 621113 7864660    1    0.13     0.76        0   0.09     0.02
## 5 20.48  21.10 621188 7865279    0    0.00     0.33        0   0.33     0.33
## 6 20.43  56.44 621199 7865261    0    0.00     0.29        0   0.57     0.14
## 7 39.80  15.03 621171 7865212    1    0.00     0.20        0   0.80     0.00
## 8 20.23 103.24 621172 7865227    0    0.09     0.18        0   0.64     0.09
## 9 19.68 117.69 621165 7865330    0    0.08     0.25        0   0.42     0.25
##      x.end1  y.end1   x.end2  y.end2 x.end3  y.end3 x.end4  y.end4
## 2 620500.6 7864660 621725.4 7864660 621113 7865272 621113 7864048
## 5 621166.9 7865279 621209.1 7865279 621188 7865300 621188 7865258
## 6 621142.6 7865261 621255.4 7865261 621199 7865317 621199 7865205
## 7 621156.0 7865212 621186.0 7865212 621171 7865227 621171 7865197
## 8 621068.8 7865227 621275.2 7865227 621172 7865330 621172 7865124
## 9 621047.3 7865330 621282.7 7865330 621165 7865448 621165 7865212
```

After the coordinates of the end point for each potential step have been calculated, extracting the corresponding LULC along each potential step relies on code that is similar to the one provided above, except that we change the coordinates that are used when creating the line segment.

```r
#extract LULC along each path
nobs=nrow(dat6)
nsteps=4
lulc.potential=numeric()
for (j in 1:nsteps){
  var.names=paste0(c('x.end','y.end'),j)
  res=matrix(NA,nobs,50)
  for (i in 1:nobs){
    print(c(j,i))

    #create line segment
    coord=cbind(c(dat6$x.sta[i],dat6[i,var.names[1]]),
                c(dat6$y.sta[i],dat6[i,var.names[2]]))
    seg1= st_linestring(coord)

    #define coordinate system
    seg2=st_sfc(seg1,crs = prj.info)
    seg3=st_sf(seg2)

    #create buffer
    seg4=st_buffer(seg3,dist=30)
    seg5=st_transform(seg4,crs=crs(ras1))

    #extract from raster
    tmp1<- raster::extract(ras1, seg5, along = TRUE, cellnumbers = F,
                           method='simple')

    #tabulate the number of each type of LULC pixel and convert to proportion
    tmp2=table(tmp1[[1]])
    tmp3=tmp2/sum(tmp2)
    tmp4=rep(0,50)
    tmp4[as.numeric(names(tmp3))]=tmp3
    res[i,]=tmp4
  }

  #just keep values that really show up
  colnames(res)=paste0('lulc',1:ncol(res))
  sum1=apply(res,2,sum)
  ind=which(sum1==0)
  res1=res[,-ind]
  colnames(res1)=paste0(c('forest','savanna','wetland','grass','pasture'),j)
  lulc.potential=cbind(lulc.potential,res1)
}
head(round(lulc.potential,2))
```

```
##      forest1 savanna1 wetland1 grass1 pasture1 forest2 savanna2 wetland2 grass2
## [1,]    0.02     0.53        0   0.34     0.11    0.24     0.67        0   0.09
## [2,]    0.00     0.00        0   0.75     0.25    0.00     0.00        0   0.25
## [3,]    0.00     0.17        0   0.67     0.17    0.00     0.67        0   0.00
## [4,]    0.00     0.20        0   0.80     0.00    0.00     0.40        0   0.60
## [5,]    0.00     0.09        0   0.91     0.00    0.00     0.80        0   0.20
## [6,]    0.08     0.08        0   0.75     0.08    0.09     0.09        0   0.09
##      pasture2 forest3 savanna3 wetland3 grass3 pasture3 forest4 savanna4
## [1,]     0.00    0.10     0.63        0   0.27     0.00    0.43     0.55
```

```
## [2,]     0.75    0.00     0.00        0   0.50     0.50    0.00     0.25
## [3,]     0.33    0.00     0.12        0   0.25     0.62    0.00     0.62
## [4,]     0.00    0.00     0.20        0   0.80     0.00    0.00     0.40
## [5,]     0.00    0.07     0.14        0   0.50     0.29    0.00     0.54
## [6,]     0.73    0.45     0.36        0   0.09     0.09    0.09     0.00
##      wetland4 grass4 pasture4
## [1,]        0   0.02     0.00
## [2,]        0   0.50     0.25
## [3,]        0   0.25     0.12
## [4,]        0   0.60     0.00
## [5,]        0   0.46     0.00
## [6,]        0   0.82     0.09
```

We combine these datasets to generate the data that will ultimately be used to fit the tSSF model.

```
#combine datasets
dat7=cbind(dat6,lulc.potential)

#remove unnecessary columns
ind=which(colnames(dat7)%in%c('x.sta','y.sta',
                              'x.end1','y.end1',
                              'x.end2','y.end2',
                              'x.end3','y.end3',
                              'x.end4','y.end4'))
dat8=dat7[,-ind]

#export tSSF data
setwd('U:\\timemod\\tutorials\\prep data tutorial')
write.csv(dat8,'tSSF data.csv',row.names=F)
```

### Time model tutorial

Denis Valle

March 2023

#### Goal

The goal of the time model is to understand how different factors influence the time it takes for an animal to traverse the landscape. Some of these factors might be associated with the animal (e.g., sex, size, and age), the landscape (e.g., amount of vegetation or water), or even more general environmental variables (e.g., temperature). It is important to note that there might be multiple reasons why an animal might spend more time on landscapes with certain characteristics. For example, these characteristics might make it more difficult for the animal to move through the landscape (i.e., resistance). Alternatively, these characteristics might make this patch of landscape ideal foraging, hiding, or resting habitat for the animal, also resulting in slower movements. On the other hand, animals might move faster through places with less resources, with short/open vegetation, and/or with higher perceived risk.

#### Data

The data come from a giant anteater monitored in the Brazilian Pantanal region. Each row in our data set consists of the information regarding a single step given by this individual:

- `time1`: the time taken for this step (minutes);
- `dist1`: the distance traversed (meters) during this step;
- `Miss`: a binary variable indicating if one or more GPS fixes were missed (=1) or not (=0) for this step. This variable was determined based on the `time1` variable. For example, if the tracking device is programmed to obtain GPS fixes every 20 minutes but the time interval between two successful GPS fixes was equal to 60 minutes, this means that two fixes (one at 20 minutes and the other at 40 minutes) were missing;
- `forest0`: proportion of forest in the area surrounding each step;
- `savanna0`: proportion of savanna in the area surrounding each step;
- `wetland0`: proportion of wetland in the area surrounding each step;
- `grass0`: proportion of grassland in the area surrounding each step;
- `pasture0`: proportion of pasture in the area surrounding each step.

It is important to note that the proportion of each land-use/land-cover (LULC) in the area surrounding each step was calculated by creating a 30-m buffer and determining the proportion of pixels associated with each LULC class. Furthermore, because parameters will be unidentifiable if all LULC classes are included, we remove grasslands/pastures from this data set and use these as the baseline LULC category.

```
rm(list=ls())
set.seed(2)

#import data
```

```
setwd('U:\\timemod\\tutorials\\prep data tutorial')
dat1=read.csv('timemod data1.csv')
head(round(dat1,2))
```

```
##   time1  dist1 Miss forest0 savanna0 wetland0 grass0 pasture0
## 1 59.92 612.38    1    0.13     0.76        0   0.09     0.02
## 2 20.48  21.10    0    0.00     0.33        0   0.33     0.33
## 3 20.43  56.44    0    0.00     0.29        0   0.57     0.14
## 4 39.80  15.03    1    0.00     0.20        0   0.80     0.00
## 5 20.23 103.24    0    0.09     0.18        0   0.64     0.09
## 6 19.68 117.69    0    0.08     0.25        0   0.42     0.25
```

These data are then organized as a list to be provided to JAGS using the code below:

```
#prepare data for JAGS
dat.jags=list(delta.time=dat1$time1,
          forest=dat1$forest0,
          savanna=dat1$savanna0,
          wetland=dat1$wetland0,
          dist=dat1$dist1,
          nobs=nrow(dat1),
          Miss=dat1$Miss)
```

#### Likelihood and priors

Recall that the likelihood for the time model is given by:

$$\Delta t_i \sim Gamma(a_i, b_i)$$

For this example, we assume that the mean of this gamma distribution is given by:

$$\mu_i = \frac{a_i}{b_i} = D_i \times exp(\beta_0 + \beta_1 Forest_i + \beta_2 Savanna_i + \beta_3 Wetland_i)$$

where $D_i$ is the distance traversed by the animal during time step $i$. Notice that we can solve this expression for $a_i$, yielding:

$$a_i = \mu_i \times b_i$$

We also allow the precision parameter $b_i$ to depend on $Miss_i$ (a binary variable equal to 1 if one or more GPS fixes were missed in time step $i$) through the function:

$$b_i = exp(\gamma_0 + \gamma_1 Miss_i)$$

The idea here is that missed GPS fixes can potentially increase the variance given that there is greater uncertainty regarding the true path of the animal. If this is true, then we expect $\gamma_1$ to be negative. If no GPS fixes were missed, then one can just drop the subscript from the precision parameter $b$ and assume that it is a constant parameter.

We finish specifying this model by assigning relatively uninformative priors to $\gamma_0$, $\gamma_1$, and $\beta_0$. On the other hand, we use priors for $\beta_1, \beta_2$ and $\beta_3$ that tend to shrink these regression parameters to zero. As a result, our analysis is more likely to be conservative (i.e., more likely to not detect an effect when it is present rather than detect an effect when it is not present).

#### JAGS model

The model described by the equations provided above is specified in a separate script entitled "resist_avg_jags.R". The content of this script is shown below:

```
#content of "resist_avg_jags.R" file

model{
  for(i in 1:nobs){
    #mean of gamma distribution
    mu[i] <- dist[i]*exp(b0+b1*forest[i]+b2*savanna[i]+b3*wetland[i])
    #factors influencing the dispersion parameter b
    b[i] <- exp(g0+g1*Miss[i])
    #calculate the corresponding a[i] parameter
    a[i] <- mu[i]*b[i]
    #likelihood
    delta.time[i] ~ dgamma(a[i],b[i])
  }

  #priors
  b0 ~ dnorm(0,0.01)
  b1 ~ dnorm(0,1)
  b2 ~ dnorm(0,1)
  b3 ~ dnorm(0,1)
  g0 ~ dnorm(0,0.01)
  g1 ~ dnorm(0,0.01)
}
```

#### Running JAGS

The regression parameters that determine mean time consist of:

- b0: intercept;
- b1: slope associated with proportion of forest;
- b2: slope associated with proportion of savanna; and
- b3: slope associated with proportion of wetland.

The parameters that govern the precision of the gamma distribution consist of:

- g0: intercept for precision; and
- g1: slope associated with missed GPS fixes.

Here we specify the settings for JAGS and finally we fit the model. Note that, to fit this model, the user will need to have installed the software JAGS, available at https://sourceforge.net/projects/mcmc-jags/files/, as this software does not come with R. We rely on the R package "jagsUI" to enable R to communicate with JAGS.

```
#jags settings and stuff
params=c('b0','b1','b2','b3','g0','g1')  #parameters to be monitored
n.iter <- 5000                #number of iterations per chain
n.thin <- 10                  #how to thin MCMC results
n.burnin <- n.iter/2          #number of iterations to discard as burn-in
```

```
n.chains <- 3                #number of MCMC chains

#run JAGS
setwd('U:\\timemod\\tutorials\\timemod tutorial')
library(jagsUI)
mod.results = jags(model.file = 'resist_avg_jags.R',
                   parameters.to.save = params, data = dat.jags,
                   n.chains = n.chains, n.burnin = n.burnin, n.iter = n.iter,
                   n.thin = n.thin, DIC = F)
```

#### Assessing convergence

We assess MCMC convergence by determining if the Rhat statistic is below 1.1 for all parameters. According to the results below, our algorithm has successfully converged.

```
mod.results$Rhat
```

```
## $b0
## [1] 1.001223
##
## $b1
## [1] 1.005197
##
## $b2
## [1] 1.005542
##
## $b3
## [1] 1.001836
##
## $g0
## [1] 1.001333
##
## $g1
## [1] 1.005215
```

#### Interpreting JAGS output

We can plot the posterior distribution of the regression parameters using the code below. This plot shows positive slope coefficients for forest and savanna, indicating that this animal tends to spend more time (i.e., moves more slowly) traversing a landscape with a greater abundance of forest and savannas when compared to grasslands/pastures (i.e., the baseline land cover). On the other hand, the slope was estimated to be negative for wetlands, revealing that this animal tends to move faster in landscapes with wetlands when compared to grasslands/pastures.

```
betas=cbind(mod.results$sims.list$b0,
            mod.results$sims.list$b1,
            mod.results$sims.list$b2,
            mod.results$sims.list$b3)
colnames(betas)=paste0('b',0:3)
boxplot(betas[,-1],las=2,names=c('forest','savanna','wetland'),ylab='Slope estimates')
abline(h=0,col='grey')
```

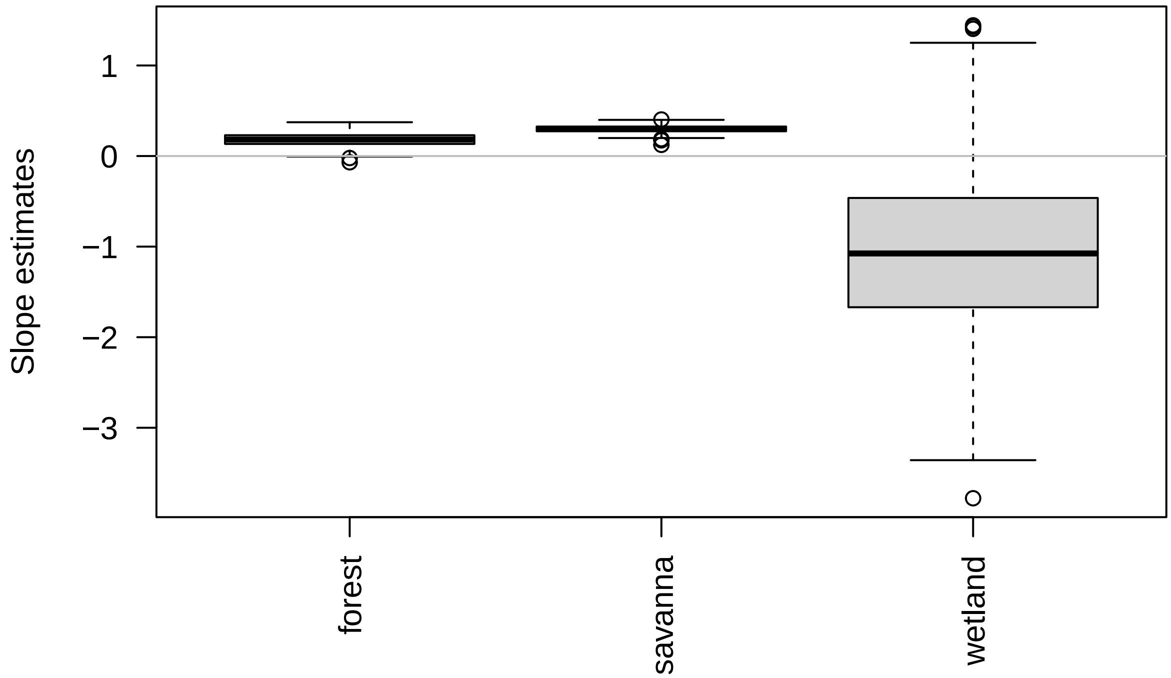


To calculate the estimated mean time taken to traverse 60 m (the median distance in the original data set) in a landscape with different proportions of savannas (for example), we rely on the following code:

```
n=100
savanna.prop=seq(from=0,to=1,length.out=n)
res=matrix(NA,n,3)
for (i in 1:n){
  linear.func=betas[,'b0']+betas[,'b2']*savanna.prop[i]
  mu=60*exp(linear.func)
  res[i,]=quantile(mu,c(0.025,0.5,0.975))
}
colnames(res)=c('lo.CI','med','hi.CI')
plot(savanna.prop,res[,'med'],ylim=range(res),type='l',ylab='Time (min)',xlab='Proportion of savanna')
lines(savanna.prop,res[,'lo.CI'],lty=2)
lines(savanna.prop,res[,'hi.CI'],lty=2)
```

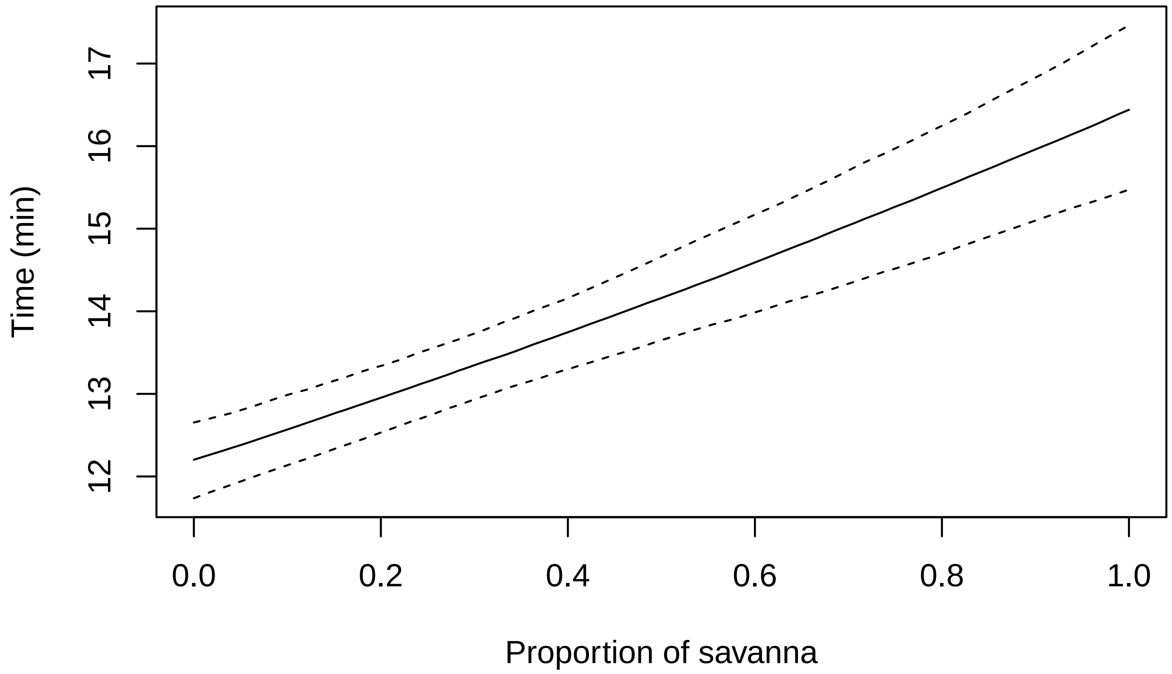


In this figure, we can see that, the higher the proportion of savanna around the animal, the slower it will move. In this case, we believe that this slower movement is due to savannas being harder to move through when compared to grasslands for giant anteaters.

We can also look at the parameters that govern the precision of the gamma distribution. We expect that missed GPS fixes can potentially increase uncertainty (i.e., decrease the precision) given that there is greater uncertainty regarding the true path of the animal. If this is true, then we expect g1 (the slope for Miss) to be negative.

The posterior distribution for these parameters reveals that, as expected, g1 is negative indicating that missed GPS fixes do indeed result in greater uncertainty.

```
gs=cbind(mod.results$sims.list$g0,mod.results$sims.list$g1)
colnames(gs)=c('g0','g1')
boxplot(gs,ylim=range(c(0,gs)))
abline(h=0,col='grey')
```

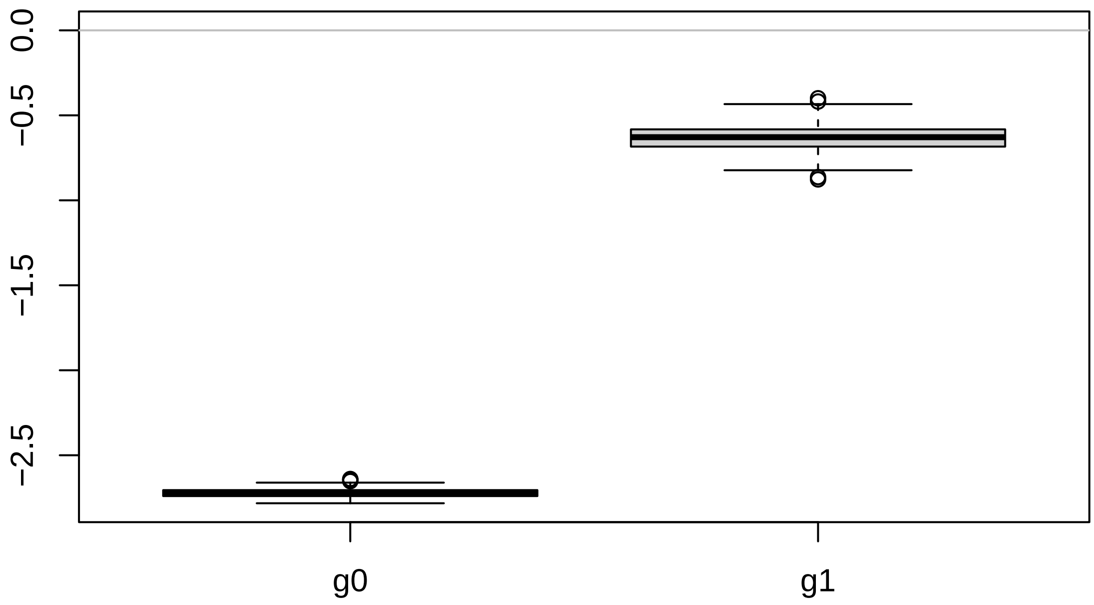


Finally, we can export our results to use for subsequent analysis.

```
setwd('U:\\timemod\\tutorials\\timemod tutorial\\results')
write.csv(betas,'betas.csv',row.names=F)
write.csv(gs,'gs.csv',row.names=F)
```

### Time-explicit Step Selection Function (tSSF) tutorial

Denis Valle

March 2023

#### Goal

The goal of the time-explicit step selection function (tSSF) is to understand how different landscape factors influence the path the animal decides to take while at the same time accounting for the feasibility of reaching each location. For instance, within a time interval of 20 minutes, it might be possible for the animal to move 60 meters through grassland to reach a particular area. However, it might be impossible for the animal to move through savannas at the same speed. We rely on the time model to determine the feasibility of reaching different areas at each moment. In other words, the time model is used to determine which areas are indeed available for the animal.

#### Initial data

The data come from a giant anteater monitored in the Brazilian Pantanal region. Each row in our data set consists of the information regarding a single step given by this individual:

- `time1` : the time taken for this step (minutes);
- `dist1` : the distance traversed (meters) during this step;
- `Miss` : a binary variable indicating if one or more GPS fixes were missed (=1) or not (=0) for this step. This variable was determined based on the `time1` variable. For example, if the tracking device is programmed to obtain GPS fixes every 20 minutes but the time interval between two successful GPS fixes was equal to 60 minutes, this means that two fixes (one at 20 minutes and the other at 40 minutes) were missing;
- `forest0` : proportion of forest in the area surrounding each step;
- `savanna0` : proportion of savanna in the area surrounding each step;
- `wetland0` : proportion of wetland in the area surrounding each step.
- `grass0` : proportion of grassland in the area surrounding each step.
- `pasture0` : proportion of pasture in the area surrounding each step.

It is important to note that the proportion of each land-use/land-cover (LULC) in the area surrounding each step was calculated by creating a 30-m buffer around the straight line that connects two consecutive locations and determining the proportion of pixels associated with each LULC class. Another important observation is that, although grasslands/pastures are present in the study region, we do not include grasslands/pastures in this data set because they are the baseline LULC category for this model.

To characterize habitat availability, we have augmented this data with the proportion of each LULC class surrounding 4 potential steps of the same length as the paired observed step:

- `forest1`,..., `forest4` : proportion of forest in the area surrounding potential steps 1,...,4;

- savanna1 ,..., savanna4 : proportion of savanna in the area surrounding potential steps 1,...,4; and
- wetland1 ,..., wetland4 : proportion of wetland in the area surrounding potential steps 1,...,4.
- grass1 ,..., grass4 : proportion of grassland in the area surrounding potential steps 1,...,4.
- pasture1 ,..., pasture4 : proportion of pasture in the area surrounding potential steps 1,...,4.

Potential steps can be selected in a variety of ways without biasing the results of the model. In this example, we used the actual step length and define four potential steps by assuming that the animal could have moved in the four cardinal directions (i.e., east, west, north, and south from where the step started). Note however that the potential steps do not even have to have the same length as the observed step for our model to work.

We start by importing these data using the code below:

```
rm(list=ls())

#get data
setwd('U:\\timemod\\tutorials\\prep data tutorial')
dat2=read.csv('tSSF data.csv')
head(round(dat2,2))
```

```
##   time1  dist1 Miss forest0 savanna0 wetland0 grass0 pasture0 forest1 savanna1
## 1 59.92 612.38    1    0.13     0.76        0   0.09     0.02    0.02     0.53
## 2 20.48  21.10    0    0.00     0.33        0   0.33     0.33    0.00     0.00
## 3 20.43  56.44    0    0.00     0.29        0   0.57     0.14    0.00     0.17
## 4 39.80  15.03    1    0.00     0.20        0   0.80     0.00    0.00     0.20
## 5 20.23 103.24    0    0.09     0.18        0   0.64     0.09    0.00     0.09
## 6 19.68 117.69    0    0.08     0.25        0   0.42     0.25    0.08     0.08
##   wetland1 grass1 pasture1 forest2 savanna2 wetland2 grass2 pasture2 forest3
## 1        0   0.34     0.11    0.24     0.67        0   0.09     0.00    0.10
## 2        0   0.75     0.25    0.00     0.00        0   0.25     0.75    0.00
## 3        0   0.67     0.17    0.00     0.67        0   0.00     0.33    0.00
## 4        0   0.80     0.00    0.00     0.40        0   0.60     0.00    0.00
## 5        0   0.91     0.00    0.00     0.80        0   0.20     0.00    0.07
## 6        0   0.75     0.08    0.09     0.09        0   0.09     0.73    0.45
##   savanna3 wetland3 grass3 pasture3 forest4 savanna4 wetland4 grass4 pasture4
## 1     0.63        0   0.27     0.00    0.43     0.55        0   0.02     0.00
## 2     0.00        0   0.50     0.50    0.00     0.25        0   0.50     0.25
## 3     0.12        0   0.25     0.62    0.00     0.62        0   0.25     0.12
## 4     0.20        0   0.80     0.00    0.00     0.40        0   0.60     0.00
## 5     0.14        0   0.50     0.29    0.00     0.54        0   0.46     0.00
## 6     0.36        0   0.09     0.09    0.09     0.00        0   0.82     0.09
```

#### Calculating the feasibility of each step

Recall that we rely on the time model to determine the feasibility of reaching different areas (i.e., to determine which areas are indeed available for the animal). For this reason, we import the parameter estimates from the time model to then calculate the feasibility of each step (i.e., calculate the gamma density associated with the realized and potential steps).

```
#get betas and gs
setwd('U:\\timemod\\tutorials\\timemod tutorial\\results')
betas=read.csv('betas.csv')
gs=read.csv('gs.csv')
```

It is important to note that, instead of explicitly writing $\beta_0 + \beta_1 \times Forest_i + \beta_2 \times Savanna_i + \beta_3 \times Wetland_i$, we rely on linear algebra to perform this calculation for each realized and potential steps. The gamma density evaluated at the realized and potential steps is stored in the matrix `prob`:

```
#get names of covariates
names.lulc=c('forest','savanna','wetland')

#calculate probabilities
prob=matrix(NA,nrow(dat2),5)
for (i in 1:nrow(dat2)){
  #get parameters
  betas1=data.matrix(betas)
  gs2=data.matrix(gs)

  #calculate probabilities for the realized and potential steps
  for (path1 in 0:4){
    names.lulc1=paste0(names.lulc,path1)
    #define design vector
    xmat=c(1,as.numeric(dat2[i,names.lulc1]))
    #calculate mean using linear algebra
    mean1=dat2$dist1[i]*exp(xmat%*%t(betas1))
    #calculate a and b parameters of gamma distribution
    b1=exp(gs2[,1]+gs2[,2]*dat2$Miss[i])
    a1=b1*mean1
    #calculate posterior median of gamma density
    prob[i,path1+1]=median(dgamma(dat2$time1[i],a1,b1))
  }
}
colnames(prob)=paste0('prob',0:4)
```

Finally, we augment our original data with these results.

```
dat=cbind(dat2,prob)
```

#### Preparing data for JAGS

We start by storing the LULC covariates for the realized steps in the matrix `xmat0` whereas the covariates for the potential steps 1,...,4 were stored in the matrices `xmat1`,...,`xmat4`, respectively. Similarly, we store the feasibility of the realized steps in the vector `pmov0` whereas for the potential steps 1,...,4, this information was stored in the vectors `pmov1`,...,`pmov4`, respectively. We also create a vector `y` comprised of ones to be able to perform the "ones trick" (described below). Finally, we put all this together in a list called `dat1`.

```
#basic settings
nobs=nrow(dat)
ncov=3
y=rep(1,nobs)

dat1=list(nobs=nobs,ncov=ncov,
          xmat0=data.matrix(dat[,c('forest0','savanna0','wetland0')]),
          xmat1=data.matrix(dat[,c('forest1','savanna1','wetland1')]),
          xmat2=data.matrix(dat[,c('forest2','savanna2','wetland2')]),
```

```
        xmat3=data.matrix(dat[,c('forest3','savanna3','wetland3')]),
        xmat4=data.matrix(dat[,c('forest4','savanna4','wetland4')]),
        pmov0=dat[,'prob0'],
        pmov1=dat[,'prob1'],
        pmov2=dat[,'prob2'],
        pmov3=dat[,'prob3'],
        pmov4=dat[,'prob4'],
        y=y)
```

#### Specifying the likelihood of a conditional logistic model

The likelihood of our model is given by:

$$L(\beta) = \prod_i \frac{pmov_{0i} \times exp(\boldsymbol{xmat_{0i}^T \beta})}{pmov_{0i} \times exp(\boldsymbol{xmat_{0i}^T \beta}) + ... + pmov_{4i} \times exp(\boldsymbol{xmat_{4i}^T \beta})}$$

where $\boldsymbol{xmat_{0i}}$ is a vector that contains the covariates for the realized step $i$ whereas $\boldsymbol{xmat_{1i}}$,. . . ,$\boldsymbol{xmat_{4i}}$ contain the covariates for the matched potential steps 1,. . . ,4, respectively. Similarly, $pmov_{0i}$ is the feasibility of the realized step $i$ whereas $pmov_{1i}$,. . . ,$pmov_{4i}$ are the feasibilities for the matched potential steps 1,. . . ,4, respectively.

In JAGS, we can specify this non-standard likelihood using the so-called "ones trick". More specifically, let

$$\pi_i = \frac{pmov_{0i} \times exp(\boldsymbol{xmat_{0i}^T \beta})}{pmov_{0i} \times exp(\boldsymbol{xmat_{0i}^T \beta}) + ... + pmov_{4i} \times exp(\boldsymbol{xmat_{4i}^T \beta})}$$

If we set $y_i$ to 1 for i=1,. . . ,n, and model this variable using a Bernoulli distribution, the likelihood will be given by the desired equation:

$$\prod_i p(y_i|...) = \prod_i \pi_i^{y_i}(1-\pi_i)^{1-y_i} = \prod_i \pi_i$$

#### JAGS model

We translate the equations provided above into the JAGS model using the code below. This code is stored in a separate file called "jags_tssf.R".

```
#content of the file "jags_tssf.R"
model{
  for (i in 1:nobs){
    #calculation of pi[i]
    p0[i] <- pmov0[i]*exp(inprod(xmat0[i,],betas))
    p1[i] <- pmov1[i]*exp(inprod(xmat1[i,],betas))
    p2[i] <- pmov2[i]*exp(inprod(xmat2[i,],betas))
    p3[i] <- pmov3[i]*exp(inprod(xmat3[i,],betas))
    p4[i] <- pmov4[i]*exp(inprod(xmat4[i,],betas))
    denom[i] <- p0[i]+p1[i]+p2[i]+p3[i]+p4[i]
    pi[i] <- p0[i]/denom[i]

    #defining likelihood using the "ones-trick"
    y[i]~dbern(pi[i])
```

```
  }

  #priors for regression parameters
  for (k in 1:ncov){
    betas[k] ~ dnorm(0,0.1)
  }
}
```

#### Running JAGS and fitting the model

Here we specify the settings of our Markov-Chain Monte Carlo (MCMC) algorithm and run JAGS to fit the model:

```
#run model
set.seed(1)
library(jagsUI)

#set parameters to monitor
params=c("betas")

# MCMC settings
ni <- 5000  #number of iterations (used to be 15000)
nt <- 10    #interval to thin
nb <- 1000  #number of iterations to discard as burn-in (used to be 5000)
nc <- 3     #number of chains

setwd('U:\\timemod\\tutorials\\tssf tutorial')
mod.results=jags(model.file="jags_tssf.R",
            parameters.to.save=params,
            data=dat1,
            n.chains=nc,
            n.burnin=nb,n.iter=ni,n.thin=nt,DIC=TRUE)
```

#### Assessing model convergence

We assess MCMC convergence by determining if the Rhat statistic is below 1.1 for all parameters. According to the results below, our algorithm has successfully converged.

```
mod.results$Rhat
```

```
## $betas
## [1] 1.000857 1.002407 1.000820
##
## $deviance
## [1] 1.003085
```

#### Interpreting JAGS output

We can plot the posterior distribution of the regression parameters using the code below. These results reveals that the slope estimates for forest and savanna were negative, indicating lower preference for forests and savannas when compared to grasslands/pastures (the baseline LULC class). On the other hand, the 95% credible intervals for wetlands encompass zero, indicating that there is no discernible difference in preference between wetlands and grasslands/pastures.

```
betas=mod.results$sims.list$betas
colnames(betas)=names.lulc
boxplot(betas,las=2,ylab='Slope estimates')
abline(h=0,col='grey')
```

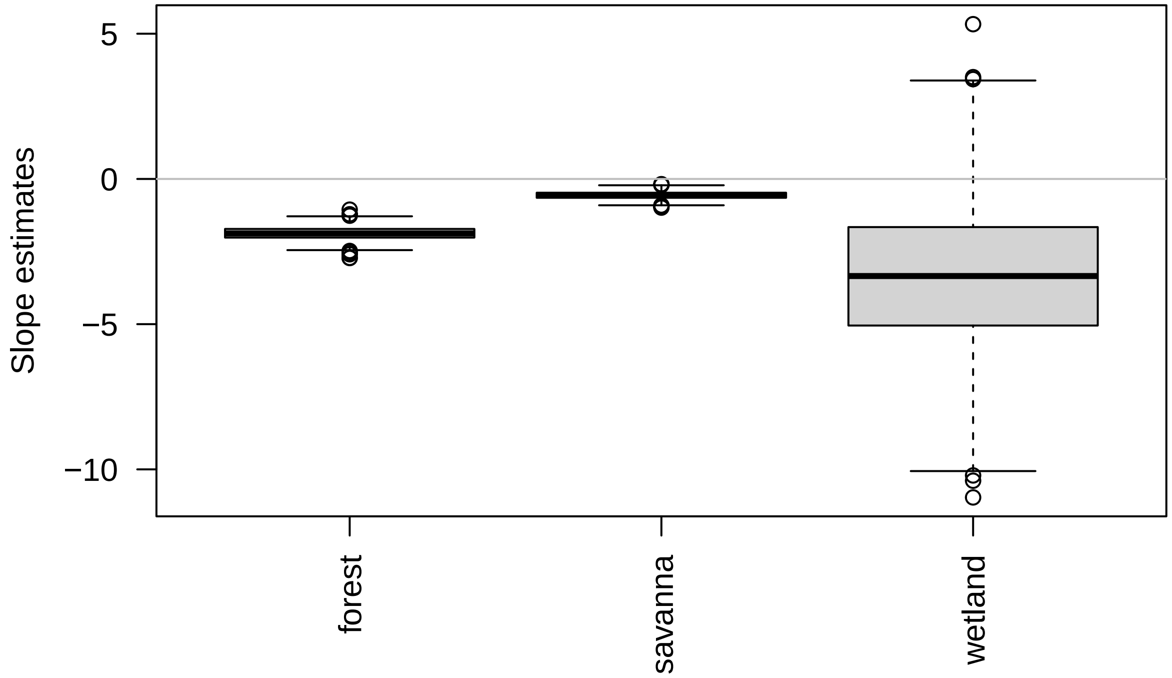


```
#calculate 95% credible intervals
apply(betas,2,quantile,c(0.025,0.975))
```

```
##           forest    savanna   wetland
## 2.5%  -2.287143 -0.8116625 -8.238711
## 97.5% -1.418803 -0.3204696  1.817181
```
